# Structural details of amyloid beta oligomers in complex with human prion protein as revealed by solid-state MAS NMR spectroscopy

**DOI:** 10.1101/2020.06.22.164574

**Authors:** Anna S. König, Nadine S. Rösener, Lothar Gremer, Daniel Flender, Wolfgang Hoyer, Philipp Neudecker, Dieter Willbold, Henrike Heise

**Affiliations:** Institute of Biological Information Processing (IBI-7: Structural Biochemistry) and JuStruct: Jülich Center for Structural Biology, Forschungszentrum Jülich, Wilhelm-Johnen-Straße, 52428 Jülich, Germany; Physikalische Biologie, Heinrich-Heine-Universität Düsseldorf, Universitätsstraße 1, 40225 Düsseldorf, Germany

## Abstract

Human PrP (huPrP) is a high-affinity receptor for oligomeric Aβ. Synthetic oligomeric Aβ species are known to be heterogeneous, dynamic and transient, rendering their structural investigation particularly challenging. Here, we used huPrP to preserve Aβ oligomers by co-precipitating them into large hetero-assemblies to investigate the conformation of Aβ(1-42) oligomers and huPrP in the complex by solid-state MAS NMR spectroscopy. The disordered N-terminal region of huPrP becomes immobilized in the complex and therefore visible in dipolar spectra without adopting chemical shifts characteristic of a regular secondary structure. Most of the well-defined C-terminal part of huPrP is part of the rigid complex, and solid-state NMR spectra suggest a loss in regular secondary structure in the last two α-helices. For Aβ(1-42) oligomers in complex with huPrP, secondary chemical shifts reveal a substantial β-strand content. Importantly, not all Aβ(1-42) molecules within the complex have identical conformations. Comparison with the chemical shifts of synthetic Aβ fibrils suggests that the Aβ oligomer preparation represents a heterogeneous mixture of β-strand-rich assemblies, of which some have the potential to evolve and elongate into different fibril polymorphs, reflecting a general propensity of Aβ to adopt variable β-structure conformers.

## Introduction

Alzheimer’s disease (AD) accounts for an estimated 60 to 80 % of all types of dementia^1^. One of the hallmarks of AD is the formation of amyloid plaques, which consist mainly of amyloid β (Aβ) peptides comprising 39 to 43 residues^2^. Aβ is produced by cleavage of the amyloid precursor protein (APP) by β- and γ-secretases^3^. Of the two most abundant species Aβ(1-40) and Aβ(1-42), the latter is more prone to aggregation and its aggregates are more toxic^3^. Small to moderately sized Aβ oligomers (Aβ_oligos_) have been identified as the most neurotoxic factor in the pathogenesis of AD, whereas large fibrils are known to be the main component of insoluble plaques^4^. Detailed structural information on Aβ(1-42)_oligo_ is still missing due to their transient nature and probably also due to high structural variability between different or within the same oligomer preparations. Heterogeneity and dynamic behavior concerning sizes and conformations is a major challenge to structural studies of oligomers^5^. This challenge has previously been met by freeze-trapping^6,7^, by applying special aggregation conditions^8^, by the addition of stabilizing compounds^9^ or antibodies^10^, or by engineered mutagenesis^11^. Here, we used the recombinant human prion protein, in its native PrP^C^ conformation to trap Aβ oligomers by co-precipitating them into large heteroassemblies, in which the growth of Aβ oligomers is prevented, as demonstrated by long-term solid-state NMR measurements over 11 months.

PrP^C^ is a high-affinity cell-surface receptor for Aβ_oligo_^12,13^ as well as for fibrillar Aβ^14–16^. It has been suggested that binding of Aβ_oligo_ to membrane-anchored PrP^C^ mediates Aβ toxicity during AD by mediating synapse damage^17^ and the blockade of long-term potentiation by Aβ_oligo_^12,18^ via activation of Fyn-kinase pathways^19,20^, but this has also been questioned^21–24^. It has also been described that soluble PrP^25^ as well as its N-terminal fragment PrP(23-111)^26,27^ have a protective role by inhibiting Aβ fibrillation and formation of Aβ_oligo_.

Several in vitro studies on the Aβ-PrP interaction suggest that Aβ_oligos_ bind at two Lys-rich parts (residues 23 to 27 and ≈ 95 to 110) on PrP^28–33^, but an additional involvement of the C-terminus of PrP has also been suggested^14^. A structural study of insoluble PrP^C^-Aβ_oligo_ complexes described this as a “hydrogel”, in which the Aβ oligomers were rigid, while PrP still has high molecular mobility^34^. Additionally, this study reported a conformational change in the N-terminus of PrP^C^ upon complexation with Aβ_oligo_. We recently demonstrated that Aβ_oligo_ forms large hetero-assemblies with either full-length (huPrP(23-230)) or C-terminally truncated (huPrP(23-144)) membrane-anchorless monomeric PrP^33^. These assemblies have a size of a few micrometers as determined by dynamic light scattering and show cloud-like morphologies as seen by atomic force microscopy^33^. The Aβ:huPrP stoichiometry of the hetero-assemblies depends on the PrP concentration added to Aβ_oligo_ and reaches a value of 4:1 (monomer ratio) in the presence of an excess of either huPrP(23-144) or huPrP(23-230)^33^. In all these in vitro preparations Aβ oligomers and early stage protofibrils are stabilized and prevented from elongation by PrP, which has been shown to preferentially bind to fast growing fibril and oligomer ends^15^.

Here we exploit this stabilizing effect in an NMR study on different samples of Aβ_oligo_ complexed by huPrP. Isotope labeling of either huPrP or Aβ allowed us to characterize both components of the complex separately. While the N-terminal region of huPrP in the complex remains largely devoid of secondary structure and still undergoes fast backbone conformational averaging on the μs to ms time scale, Aβ_oligos_ exhibit a high degree of β-strand conformation. While these Aβ_oligos_ are highly heterogeneous, solid-state NMR spectra reveal similarities with corresponding spectra of all fibril polymorphs published so far^35–37^.

## Results

### The N-terminal construct huPrP(23-144) is disordered in solution at mildly acidic and neutral pH

The solution structure of huPrP(23-230) had originally been determined in acetate buffer at an acidic pH of 4.5 and 20 °C^38^, whereas the huPrP-Aβ(1-42)_oligo_ complex samples for solid-state NMR were prepared at a pH value close to neutral. As a basis for studying the interaction between huPrP and Aβ_oligo_ we therefore first investigated free huPrP(23-144) by NMR spectroscopy in solution at different pH values ranging from 4.5 to 7.0 and at a temperature of 5.0 °C, which is closer to the temperature used for the solid-state NMR experiments. As reported previously, the chemical shifts of the N-terminal amino acid residues 23 to 124 in truncated huPrP (23-144) are almost identical to those of huPrP(23-230), whereas residues 125 to 144, which are part of the well-ordered globular domain of huPrP(23-230), are strongly affected by the truncation at position 144 ^33^.

We obtained almost complete sequence-specific ^1^H, ^13^C, and ^15^N backbone resonance assignments for huPrP(23-144) at pH values of 4.5 and 7.0 and a temperature of 5.0 °C using a combination of HNCO, HNCACB, and BEST-TROSY-(H)N(COCA)NH triple-resonance experiments (**Supplementary Fig. 1**). The assigned chemical shifts at pH 4.5 and pH 7.0 have been deposited with the Biological Magnetic Resonance Data Bank (BMRB) under accession codes 28115 and 28116, respectively.

As expected, side-chain titration in this pH range causes significant chemical shift changes for all 7 histidine residues and for residues next to histidine. Other than that, the chemical shifts at pH 4.5 and pH 7.0 are very similar to each other and very close to random coil shifts^39^. Quantitative analysis reveals that the Random Coil Index (RCI) order parameters^40^S_RCI_^2^, which are a measure of how different the backbone chemical shifts are from those of a disordered random coil on a scale of 0 (typical for a random coil) to 1 (typical for a well-ordered backbone conformation), are consistently below ≈ 0.6 (**Supplementary Fig. 2**). This demonstrates conclusively that free huPrP(23-144) in solution at neutral and mildly acidic pH is highly disordered and devoid of any stable secondary structure.

### The flexible N-terminus of huPrP becomes immobilized but remains almost devoid of regular secondary structure upon binding to Aβ_oligo_

Formation of high molecular weight assemblies was analyzed by sucrose density gradient ultracentrifugation (DGC) and subsequent SDS-PAGE, and RP-HPLC^33^ (**Supplementary Fig. 3**). This indicates that the potential PrP binding sites on Aβ_oligo_ are not yet saturated with huPrP(23-144). As previously described^33^, a molar ratio of Aβ:PrP of 4:1 is obtained in the precipitate if huPrP is added in excess.

To probe the flexibility of the N-terminal construct huPrP(23-144) in the complex, we recorded a ^1^H-^13^C Insensitive nuclei enhanced by polarization transfer (INEPT)-NMR spectrum as well as dipolar based ^1^H-^13^C and ^1^H-^15^N CP-MAS spectra^41^. The INEPT-NMR spectrum of this sample did not show any protein signals at a temperature of ≈ 27 °C (spectrum not shown), whereas in ^1^H-^13^C and ^1^H-^15^N CP spectra obtained at ≈ 0 °C strong signals typical for all amino acid types can be seen (**Supplementary Fig. 4**). This indicates that huPrP(23-144) in complex with Aβ(1-42)_oligo_ is immobilized and does not undergo rapid isotropic reorientation as in solution.

In **Fig. 1** a typical 2D ^13^C-^13^C-correlation spectrum obtained with proton driven spin diffusion (PDSD) of huPrP(23-144)*-Aβ (* indicates that the huPrP moiety of the complex is ^13^C/^15^N labelled) is overlaid with a ^13^C-^13^C total correlation spectrum (TOCSY) of monomeric huPrP(23-144) in solution at pH 6.7. Except for some Val and Ala resonances, most of the peaks align well. This indicates that the natively unfolded N-terminus of huPrP does not undergo a major conformational rearrangement upon complex formation with Aβ_oligo_, but conformational averaging of backbone conformations is still possible on the μs to ms time scale. Due to the lack of secondary structure in the intrinsically unstructured N-terminus as well as the repetitiveness of the amino acid sequence in the octarepeats the signal overlap is so severe that sequence-specific resonance assignment for the solid-state NMR spectra was not possible.

**Fig. 1.**
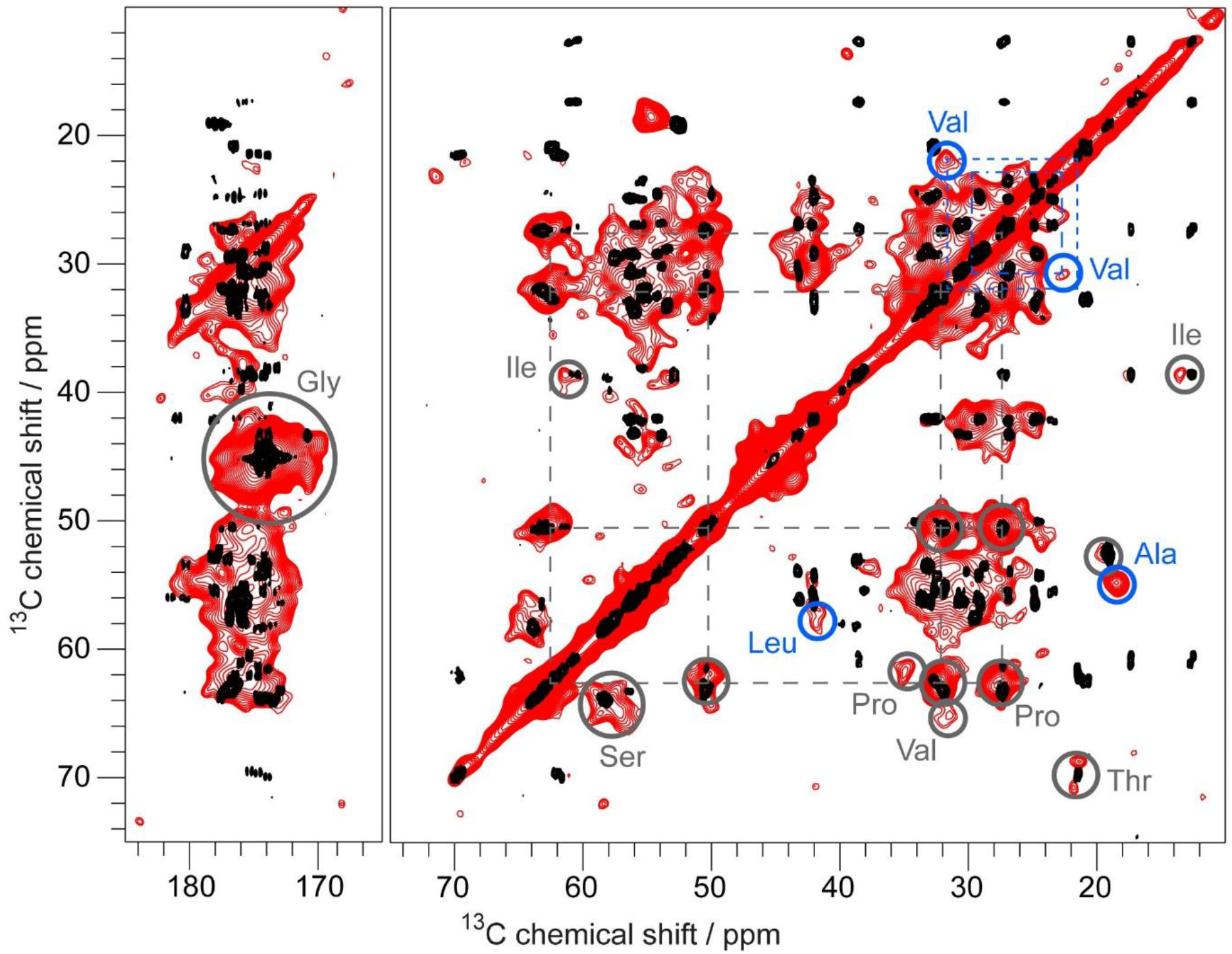
Comparison of a PDSD spectrum of huPrP(23-144)*-Aβ (* species is ^13^C, ^15^N uniformly labeled) in red with a ^13^C-^13^C TOCSY spectrum of monomeric huPrP(23-144) in black. The PDSD spectrum was recorded at a temperature of ≈ −6 °C, a spinning frequency of 11 kHz and a mixing time of 30 ms and the TOCSY spectrum at a temperature of 5 °C, at pH 6.7. Grey circles indicate some identified amino acid types, dashed lines Pro and Val connections in the PDSD spectrum. Due to broad line widths and a low signal dispersion in the PDSD spectrum several correlations overlap, especially for the residues in the octarepeat region. Nevertheless, spin systems for most of the amino acid types present in the sequence could be identified and an amino acid-type specific resonance assignment was possible. Differences between the PDSD and TOCSY spectrum are highlighted with blue circles. For an additional PDSD spectrum see **Supplementary Fig. 5**, the corresponding double-quantum single quantum correlation spectrum (DQ-SPC5) is shown in **Supplementary Fig. 6**.

While most of the resonances of huPrP(23-144) in complex with Aβ_oligo_ align well with those of huPrP(23-144) in solution, some differences can be clearly seen; in particular, some Ala, Val and Leu resonances are shifted from random coil towards α-helical secondary chemical shifts. Six out of seven Ala residues as well as both Val and Leu residues present in the sequence are located within a short stretch from residue 113 to 130, a region which starts with the so-called palindrome segment (A^113^GAAAAGA^120^) (see **Fig. 2a**). Thus, structural changes upon complexation with Aβ_oligo_ in N-terminal huPrP seem to be confined to the region between A113 and L130.

**Fig. 2.**
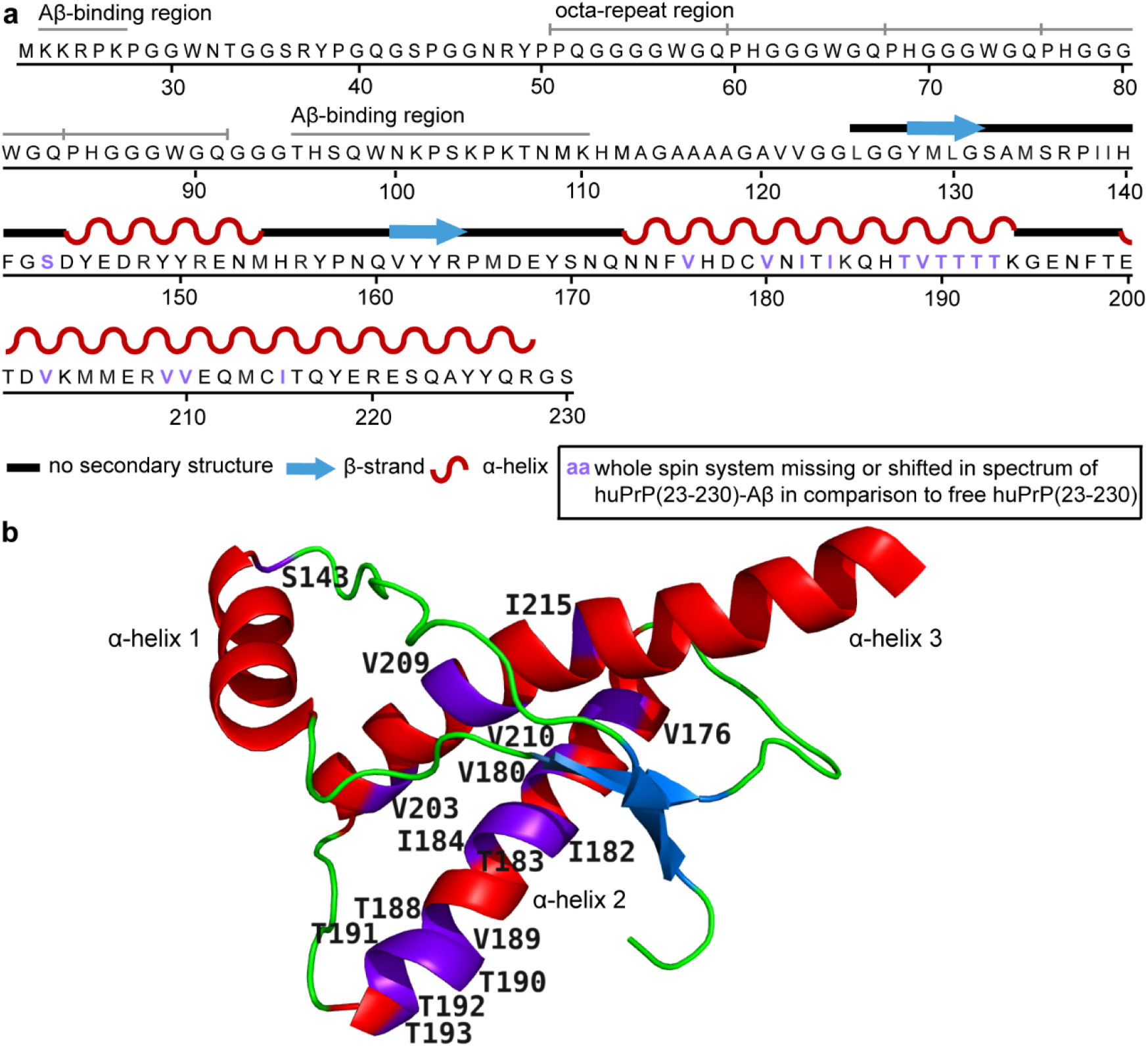
**a** Amino acid sequence of huPrP(23-230)^38^. The Aβ-binding regions K23 to K27 and T95 to K110^28–33^ and the five octarepeats are indicated above the sequence. **b** 3D structure of the natively folded prion domain (residues 125 to 228) of full-length huPrP(23-230) in solution. β-strands are colored blue, α-helices red. Picture adapted from PDB-File 1QLZ^38^. Residues whose entire spin system is missing or shifted in the PDSD spectra (**Fig. 1**) are highlighted in purple in **a** and **b**.

To exclude self-aggregation of huPrP(23-144) in the investigated samples, we also compared the chemical shifts of huPrP(23-144) fibrils^42,43^ with our correlation spectrum of huPrP(23-144)*-Aβ (**Supplementary Fig. 7**). Most of the signals observed for fibrillar huPrP(23-144) do not overlap with the signals in our huPrP(23-144)*-Aβ spectra. We therefore conclude that the conformations of huPrP(23-144) in huPrP(23-144)*-Aβ and the huPrP(23-144) fibril are very different, and the interaction with Aβ_oligo_ did not induce huPrP(23-144) fibril formation.

### The C-terminus of huPrP shows changes in α-helices 2 and 3 upon Aβ_oligo_ binding

For full-length huPrP in complex with Aβ_oligo_, huPrP(23-230)*-Aβ (see **Table 1**) no signals were detected at ≈ 30 °C in the INEPT spectrum (not shown), which indicates that not only the N-terminus, but also the C-terminus of huPrP is not highly dynamic. By contrast, a ^1^H-^13^C CP spectrum recorded at ≈ 0 °C displays the full signal set expected for a protein (**Supplementary Fig. 8**).

**Table 1:** Details of the samples used for the solid-state NMR measurements (* species is ^13^C, ^15^N uniformly labeled)

| sample | $^{13}\text{C}$, $^{15}\text{N}$ labeled species | monomer stoichiometry Aβ:huPrP (initial mixture) | monomer stoichiometry Aβ:huPrP (in complexes) |
| --- | --- | --- | --- |
| huPrP(23-144)*-Aβ | huPrP(23-144) | 6:1 | not estimated |
| huPrP(23-230)*-Aβ | huPrP(23-230) | 4:1 | not estimated |
| huPrP(23-144)-Aβ* | Aβ(1-42) | 8:1 | 8.6 : 1 |
| huPrP(23-144)<sub>exc</sub>-Aβ* | Aβ(1-42) | 2:1 | (3.7 ± 0.12) : 1 |

In **Fig. 3**, a typical 2D PDSD spectrum of the huPrP(23-230)*-Aβ complex is displayed. Again a line width of ≈ 1 ppm is observed for the ^13^C resonances, and due to the large number of resonances and the limited signal dispersion, the signal overlap is so substantial, that a sequential resonance assignment or even a quantitative analysis of residue-specific correlations was not possible. Nevertheless, a comparison with the corresponding 2D ^13^C-^13^C correlation spectrum of the N-terminal construct huPrP(23-144) in complex with oligomeric Aβ (red outline in **Fig. 3**) allows some conclusions about the structure of full-length complexed huPrP: First, almost all resonances observed in the spectrum of C-terminally truncated huPrP(23-144) appear to be also visible in the spectrum of full-length huPrP(23-230)*-Aβ (**Fig. 3**). These findings suggest that the C-terminus of full-length huPrP(23-230) does not have a major impact on the conformation of the N-terminus and its interaction with Aβ_oligo_, in line with previous results^28–30,33^. Second, the spectrum of full-length huPrP complexed by Aβ_oligo_ displays additional resonances which are absent in the spectrum of N-terminal huPrP(23-144) in complex with Aβ_oligo_. Some of the amino acid residues occurring mainly in the C-terminus (e.g. Ile, Thr and Val) give rise to cross-correlation signals that can be unambiguously identified in 2D ^13^C-^13^C cross-correlation spectra. However, for most C-terminal amino acid residues (e.g. Asp, Glu, Tyr etc., **Fig. 2a**) the 2D correlations overlap with other resonances and can therefore not be unambiguously assigned. Further, higher flexibility of the C-terminus of huPrP as compared to the N-terminus may result in reduced signal intensity for amino acid residues from those regions.

**Fig. 3.**
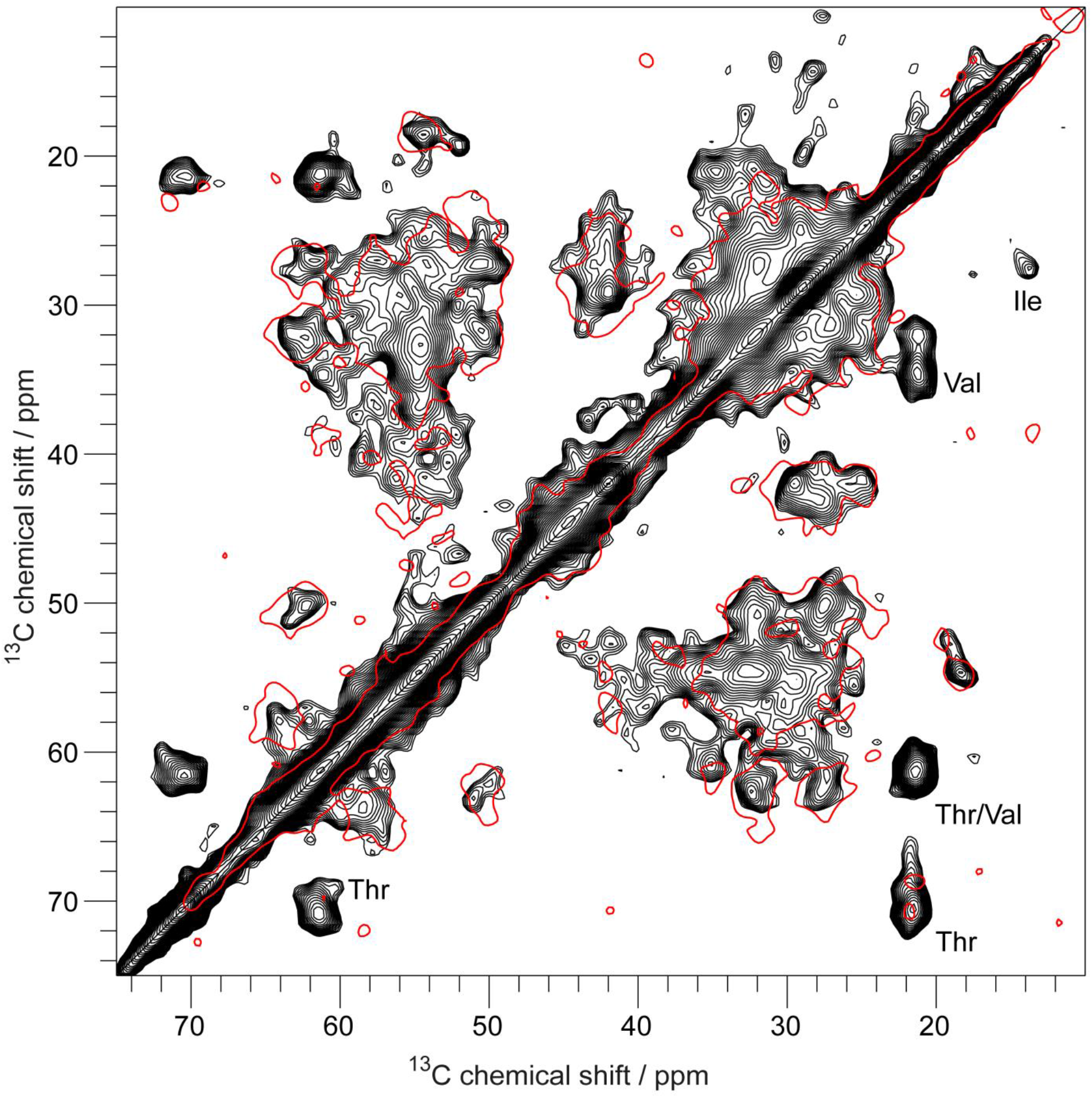
Comparison of two PDSD spectra of huPrP(23-230)*-Aβ (* species is ^13^C, ^15^N uniformly labeled), shown in black, and huPrP(23-144)*-Aβ, shown as red contour. Both spectra were recorded at a spinning frequency of 11 kHz and a mixing time of 30 ms, but the black one at a temperature of ≈ 0 °C and the red one at ≈ −6 °C.

We compared our 2D ^13^C-^13^C correlation spectrum with the expected correlations between the chemical shifts obtained experimentally for natively folded full-length huPrP in solution at pH 4.5 ^38^. While the predicted N-terminal cross peaks (residues 23 to 124) superimpose well with the spectrum of huPrP(23-230)*-Aβ, some discrepancies between the experimental and the predicted spectrum are observed for the C-terminus (residues 125 to 230) (**Supplementary Fig. 9**).

In particular, correlation signals for Ile, Thr and Val in α-helical conformation from α-helices 2 and 3 in natively folded huPrP are completely missing in the experimental spectrum (**Supplementary Fig. 10**). Instead, correlation signals for Thr and Val with secondary chemical shifts indicative of β-strands that are not observed in natively folded huPrP are clearly visible in the experimental spectrum (**Supplementary Fig. 9 to 11**). This suggests that at least for a substantial fraction of the huPrP molecules within the complex some parts of a region between either V121 and I139 and/or V176 and I215 (located in α-helices 2 and 3) have undergone some structural rearrangements including β-strand formation (**Fig. 2b**).

### High β-strand content of Aβ_oligo_ in huPrP(23-144)-Aβ complexes

We also investigated the homogeneity and structural characteristics of Aβ_oligo_ using two samples containing uniformly ^13^C, ^15^N labeled Aβ_oligo_ in complex with non-labeled huPrP(23-144) in different molar ratios (**Table 1**).

INEPT spectra recorded at ≈ 20 °C of both samples are devoid of protein signals (not shown), whereas ^1^H-^13^C and ^1^H-^15^N CP spectra recorded at ≈ 0 °C display strong signals typical for all amino acid residue types (**Supplementary Fig. 12**). These findings indicate that also the Aβ molecules are rigid parts of the complex. In all 2D and 3D homonuclear ^13^C-^13^C and heteronuclear ^15^N-^13^C correlation spectra (see **Fig. 4** and **Supplementary Fig. 13 to 17**) linewidths are rather broad (0.9 ppm for ^13^C and 3.3 ppm for ^15^N) which is an indication for conformational heterogeneity of the Aβ molecules within the complex. In 2D ^13^C-^13^C correlation spectra (**Fig. 4** and **Supplementary Fig. 13**) ^13^C side chain and backbone resonances can be identified for almost every amino acid residue type in the sequence. For several amino acid residue types, the number of distinct spin systems visible in the spectra is larger than the number of amino acid residues of this type in the amino acid sequence. For example, for Ala six spin systems have been found although the sequence of Aβ(1-42) only contains four Ala residues (**Fig. 4**). This means that not all Aβ molecules within the complex experience identical environments.

**Fig. 4.**
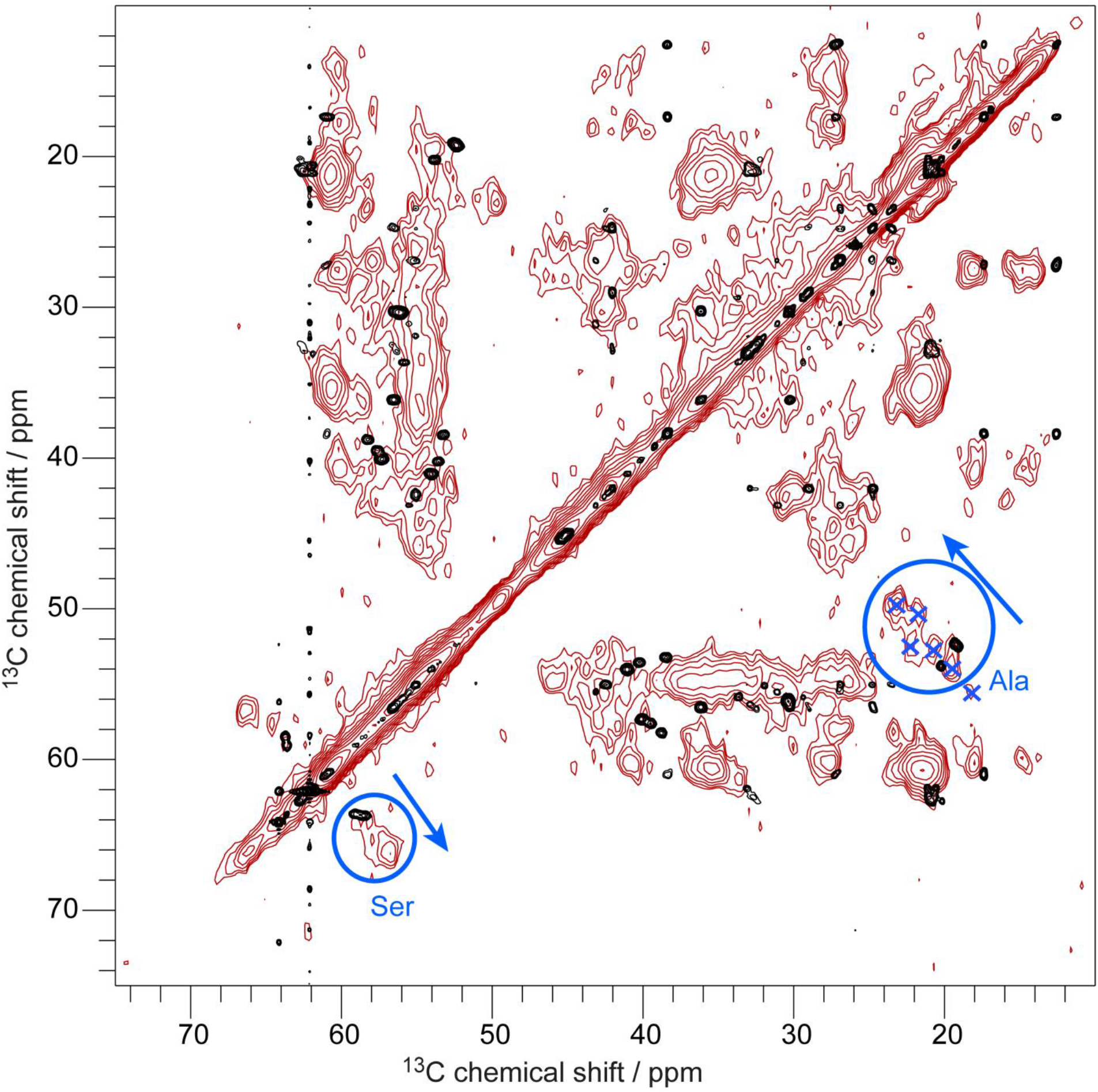
Overlay of a PDSD spectrum of huPrP(23-144)-Aβ* (* species is ^13^C, ^15^N uniformly labeled), measured at a temperature of ≈ 0 °C, a spinning frequency of 11 kHz and a mixing time of 50 ms, in red with a ^13^C-^13^C TOCSY spectrum of uniformly ^13^C, ^15^N isotope labeled Aβ monomers in solution (measured at a temperature of 5 °C and pH 7.2 in 30 mM Tris-HCl buffer) in black (the strong resonances at 62.1 ppm and 64.2 ppm with the t1 noise are from the Tris buffer). Ala and Ser C_α_-C_β_ peaks are highlighted with blue circles and six identified Ala spin systems are shown with blue crosses. Blue arrows indicate the shift changes of both residues due to a change from random coil in Aβ monomers to more β-strand like conformations in huPrP(23-144)-Aβ*.

A comparison between a solid-state NMR ^13^C-^13^C correlation spectrum of Aβ_oligo_ in complex with huPrP(23-144) and a ^13^C-^13^C TOCSY correlation spectrum of Aβ monomers in solution (**Fig. 4**) reveals strong chemical shift differences and thus indicates that the Aβ monomer building blocks in oligomers have undergone significant structural changes upon oligomerization. While all signals of the solution spectrum have chemical shifts indicative of a random coil, a strong shift to secondary chemical shifts indicative of β-strand like secondary structure is observed for almost all spin systems of Aβ_oligo_ in the spectrum of the complex. For Cα/Cβ cross correlations of Ala, Gly, Ile, Ser and Val (**Fig. 4**) in α-helical, unstructured and β-strand like conformations a quantification was possible by integration of the peak regions (see **Supplementary Fig. 18**). Hence, these residues are predominantly in a β-strand conformation, except for Gly, which is a β-strand breaker.

Due to conformational heterogeneity, inhomogeneous line broadening and substantial resonance overlap in the ^13^C-^13^C and ^15^N-^13^C spectra, a full sequential resonance assignment for Aβ_oligo_ in complex with huPrP was not possible. However, from a series of PDSD spectra with different mixing times as well as 2D and 3D NCACX and NCOCX spectra it was possible to identify some interresidual correlations and to obtain site-specific assignments for some parts of Aβ in one predominant conformation (**Supplementary Table 1**). However, it is hard to tell whether all assigned resonances belong to one type of conformer, or to different conformers.

To elucidate whether the stoichiometry of Aβ and huPrP in the heteroassemblies has an influence on the conformations of Aβ molecules, we prepared and investigated a second sample, in which huPrP(23-144) was added in excess to ^13^C, ^15^N labeled Aβ_oligo_. In this sample, all potential huPrP binding sites on Aβ_oligo_ should be occupied. Overall there is not much difference between sample huPrP(23-144)-Aβ* and huPrP(23-144)_exc_-Aβ* in a PDSD spectrum with a mixing time of 50 ms, except for minor changes (**Supplementary Fig. 19**). As there are no major structural changes upon altering the huPrP concentration, we conclude that the conformational heterogeneity is not due to unoccupied huPrP binding sites in Aβ_oligo_, but rather Aβ_oligo_ in complex with huPrP consists of inequivalent conformers and/or Aβ_oligo_ assemblies are different from each other.

## Discussion

In this study we investigated the structures and the interaction of Aβ(1-42)_oligo_ and huPrP by solid-state NMR spectroscopy (see **Fig. 5** for a schematic representation of the structural features of the huPrP-Aβ_oligo_ complex).

**Fig. 5.**
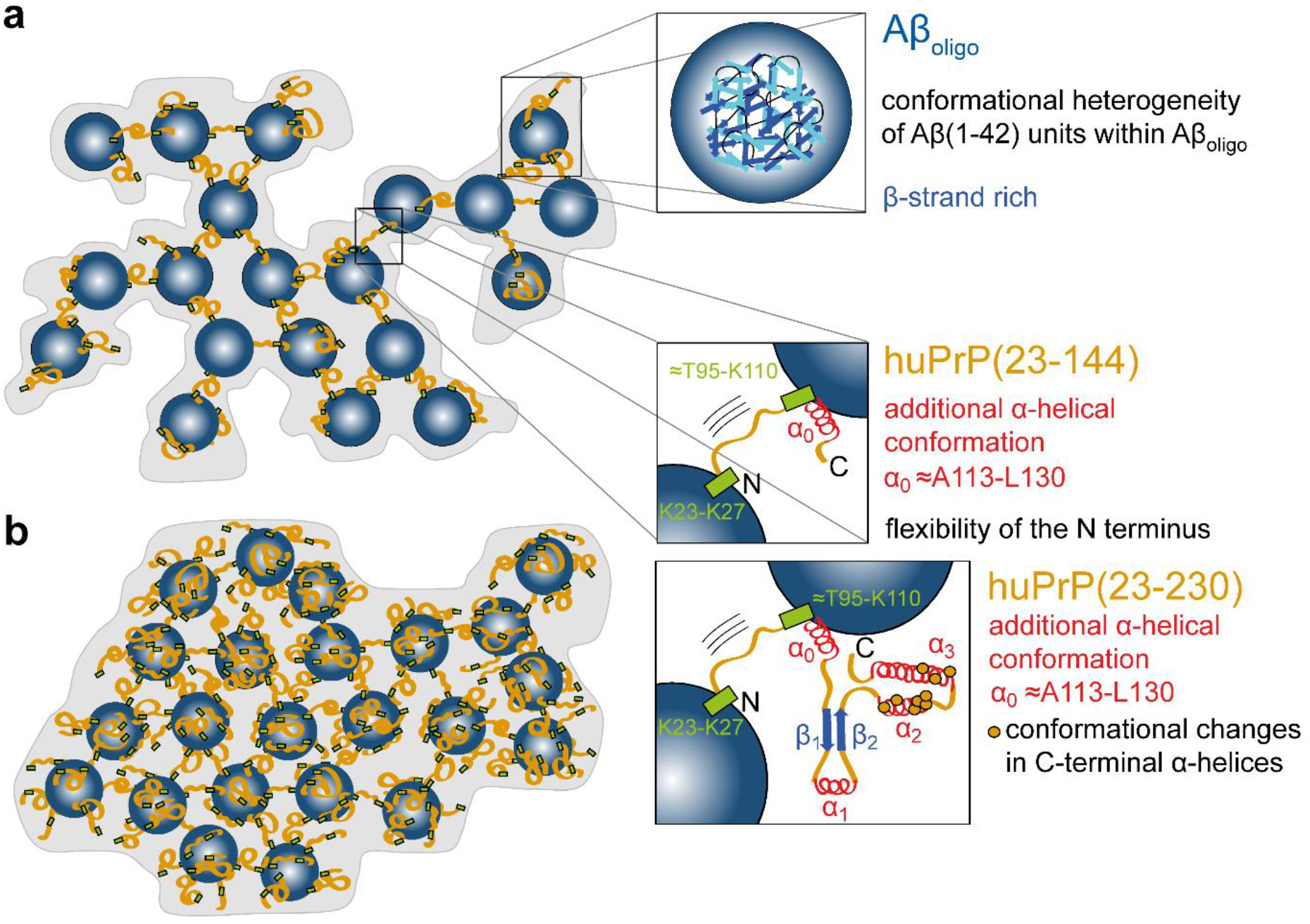
Schematic representation showing structural features of the huPrP-Aβ_oligo_ complex for **a** low huPrP content as in the sample where huPrP(23-144) is not in excess (huPrP(23-144)-Aβ*) and **b** with high huPrP content as in the sample where huPrP(23-144) was added in excess (huPrP(23-144)_exc_-Aβ*). huPrP is shown as orange lines, Aβ_oligo_ as blue spheres, α-helices are red and β-strands blue. Binding regions at huPrP are shown as light green boxes, conformational changes in the C-terminus of huPrP as orange dots. Picture adapted from Rösener et al.^33^.

The N-terminal region of huPrP is rigid but has no regular secondary structure in the complex with Aβ_oligo_. This is the case for both huPrP constructs, the full-length construct huPrP(23-230) and the N-terminal fragment huPrP(23-144). Minor structural changes to more α-helical like secondary structure are restricted to a region between A113 and L130, including the palindrome region. This palindrome region, known as the ‘hydrophobic core’, is highly conserved, and the AGAAAAGA region is highly amyloidogenic^44^. The palindrome segment has previously been suggested to be required for the attainment of the PrP^Sc^ conformation and to facilitate the proper association of PrP^Sc^ with PrP^C^ to enable prion propagation^45^. Trapping the ‘hydrophobic core’ by binding to Aβ(1-42)_oligo_ might explain the Aβ(1-42)-oligomer induced inhibition of prion propagation proposed by Sarell et al.^46^. In a recent study of a hydrogel-termed complex of full-length huPrP and Aβ_oligo_^34^ the formation of two additional α-helices, one in the octarepeat region (residues 51 to 91) and one in the palindrome segment (A^113^GAAAAGA^120^), was postulated from the observation that chemical shifts observed for Gly and Ala are predominantly α-helical in their spectra^34^. Our results support the formation of the latter α-helix in the complex with full-length huPrP. Chemical shifts of Gly residues as well as all other N-terminal residues are predominantly random coil-like in the spectra (see **Fig. 1**), suggesting that the octarepeat region does not undergo major structural rearrangements upon complex formation.

For full-length huPrP in complex with Aβ_oligo_ we observed some changes for Thr and Val residues from α-helical to random coil secondary chemical shifts compared to well-folded monomeric huPrP in solution^38^. huPrP was previously found to undergo a liquid-liquid phase separation in PBS buffer at pH 7.4 prior to complexation with Aβ_oligo_^34^. For pure liquid huPrP as well as for huPrP in a hydrogel with Aβ_oligo_ chemical shift changes from α-helical to random coil values were described for Thr residues, which are mainly located in α-helices 2 and 3 ^34^. This observation was attributed to a loss of secondary structure during liquid-liquid phase separation and is still visible in the complex^34^. This fits to our observation that no cross correlation signals typical for Thr in α-helical conformation (from α-helix 2 and 3 in natively folded huPrP) were observed in our spectra.

Aβ_oligo_ in complex with huPrP consists of non-identical Aβ conformers. This is not surprising given the fact that the sample huPrP(23-144)-Aβ* contains eight times more Aβ (monomer equivalent) than huPrP(23-144) molecules. Not every monomer within the oligomer (containing 23 monomer units on average^47^) might be able to bind to huPrP(23-144) in the same way and have therefore the same conformation^33^, as described above. These non-identical conformers can have different origins: (i) different types of monomers within the oligomer, because not every monomer can bind to huPrP (Aβ-huPrP vs. Aβ-Aβ interactions);(ii) polymorphism within the oligomer independent of the binding to huPrP (iii) polymorphism between different oligomers; or (iv) a combination thereof.

The secondary structure of Aβ_oligo_ in complex with huPrP shows a high degree of β-strand content. This could be due to fibrils that evolved over time. However, we took care not to obtain fibrils in our samples during preparation and there were no major shift changes in the CP and PDSD spectra in the following eleven months. This indicates that Aβ_oligos_ already contain Aβ monomer units that have at least in part the same secondary structure as in fibrils or protofibrils. This phenomenon has already been observed in early stage Aβ oligomers^6,7^. Assuming that the Aβ_oligo_ preparation yielded a heterogeneous collection of “small fibrils”, of which most if not all were obviously elongation incompetent, when trapped by adding huPrP, one would expect that the solid-state NMR resonances of Aβ_oligo_ in complex with huPrP are the sum of the resonances of different fibril conformations together with resonances from Aβ units that experience different environments due to edge effects and/or huPrP interaction. To assess the structural similarity of Aβ_oligo_ with fibrils and protofibrils, we superimposed all available resonances from three different Aβ(1-42) fibril types^35–37^ and one artificial protofibril^11^ with the PDSD spectrum of Aβ_oligo_ in complex with huPrP (**Fig. 6**). Nearly all predicted correlations from these different protofibril and fibril types are represented by correlation peaks in our oligomer spectra. These findings suggest that the Aβ_oligo_ preparation represents a heterogeneous mixture of β-strand-rich assemblies, of which some may have the potential to evolve into the different fibril types. The conformational heterogeneity of Aβ_oligo_ is closely related to the polymorphism of Aβ fibrils and reflects the general propensity of Aβ to adopt variable β-structure conformers.

**Fig. 6.**
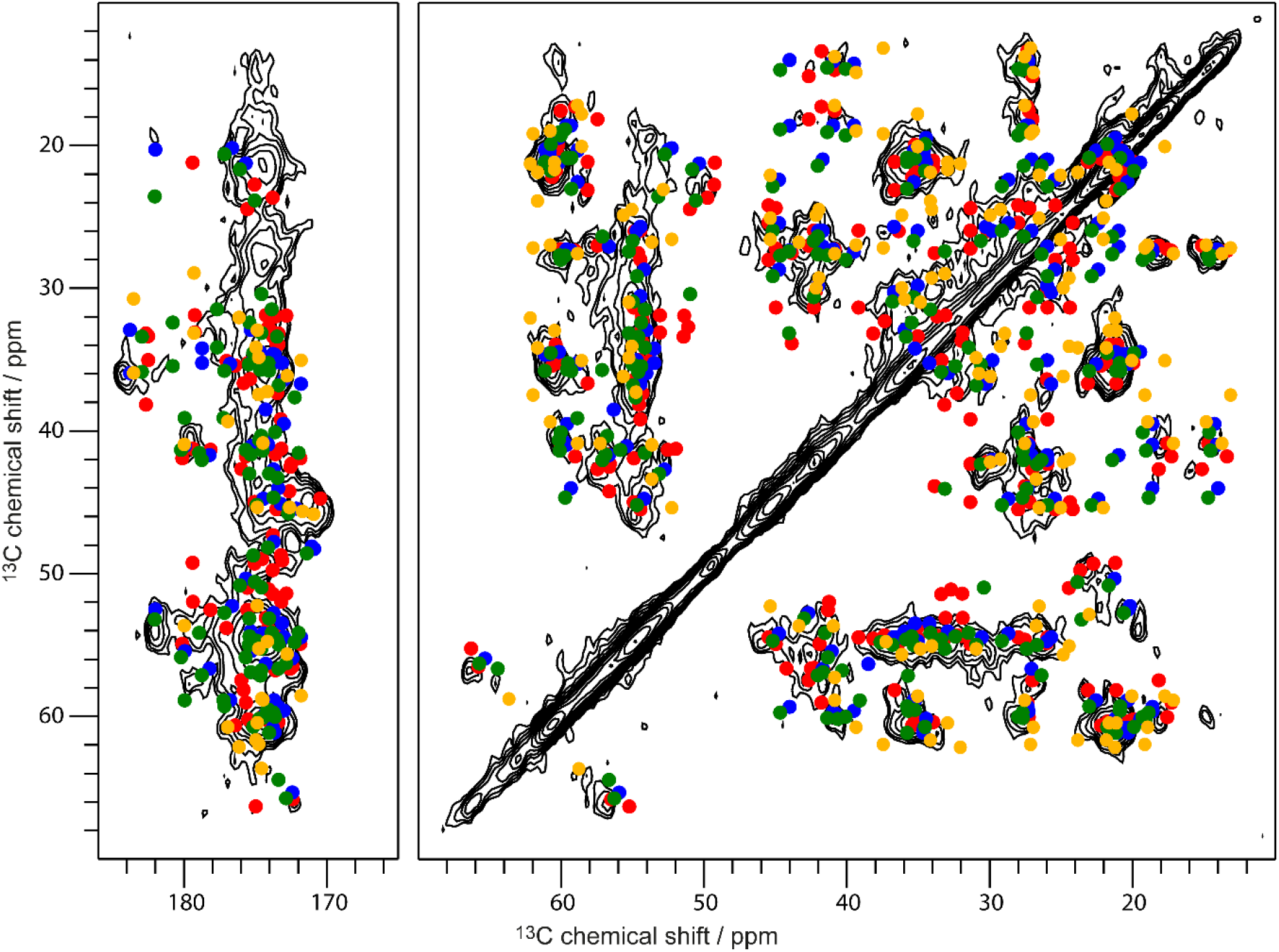
A PDSD spectrum (measured at a temperature of ≈ 0 °C, a spinning frequency of 11 kHz and a mixing time of 50 ms, same spectrum as in **Fig. 4**) of huPrP(23-144)-Aβ* (* species is ^13^C, ^15^N uniformly labeled) in comparison with predicted cross correlations (up to two bonds) for three different fibril types which are obtained at pH values of 2 (red)^37^ or 7.4 (green ref. ^36^ and blue ref. ^35^) and an artificial protofibril (yellow)^11^. Separate overlays of this PDSD spectrum with spectra of these fibrils are shown in **Supplementary Fig. 20 to 23**.

The propensity of huPrP to efficiently bind to Aβ_oligo_ and to “freeze” them in a non-dynamic and non-elongating state allowed us to investigate the conformers of Aβ_oligo_ and the huPrP moiety by NMR over several months without noticeable changes in the sample. It is tempting to speculate whether this property of huPrP is a coincidence, or whether it is part of the long-sought function of PrP. Regardless of whether PrP inhibits elongation of Aβ oligomers and fibrils or whether PrP is a mediator of cytotoxicity of Aβ_oligos_, substances that compete with PrP for Aβ_oligo_ binding and which thus can do the same job without the potential of mediating cytotoxicity may be of high therapeutic potential.

## Material and Methods

### Proteins

#### Aβ

For preparation of NMR samples with unlabeled Aβ, synthetic Aβ(1-42) obtained from Bachem AG was used. (For preparation of stocks see below.) Uniformly ^13^C, ^15^N labeled Aβ(1-42) was purchased from Isoloid GmbH.

#### huPrP

The purification of recombinant full-length huPrP(23-230) and C-terminally truncated huPrP(23-144) either unlabeled or uniformly ^13^C, ^15^N labeled was performed as described previously^33^.

### Preparation of Aβ(1-42) stocks

Synthetic unlabeled Aβ(1-42) (Bachem AG, 1 mg aliquot) was incubated with 700 μl hexafluoro-2-propanol (HFIP) overnight and divided into 108 μg doses in LoBind reaction tubes (Eppendorf AG). Samples were lyophilized in a rotational vacuum concentrator system connected to a cold trap (both Martin Christ Gefriertrocknungsanlagen GmbH). The lyophilizates were stored at room temperature and protected from light.

### Preparation of high molecular weight hetero-assemblies from amyloid β oligomers and different human prion protein constructs in different molecular ratios

For sample preparation, Aβ(1-42) lyophilisates (either uniformly ^13^C, ^15^N labeled or unlabeled) were dissolved in 30 mM Tris-HCl buffer, pH 7.4, yielding Aβ(1-42) concentrations of 160 – 300 μM. After 2 h of incubation at 22 °C and 600 rpm shaking to obtain Aβ_oligo_, either huPrP(23-144) or huPrP(23-230) was added to yield concentrations of 40 – 80 μM within the initial mixture leading to the molar ratios mentioned in **Table 1**. The addition of huPrP resulted in immediate sedimentation of the complex^33^.

After addition of 0.03 % of sodium azide and incubation for 30 min the samples were centrifuged for two to five minutes at 16,100 x *g*, and the supernatant was removed. The sediment was washed twice with up to 2 ml of 30 mM Tris-HCl buffer, 0.03 % sodium azide, pH 7.4. After removal of the supernatant the samples were transferred into 3.2 mm MAS rotors with a Hamilton syringe and centrifuged. In total, four different samples were prepared in which either huPrP or Aβ(1-42) was uniformly ^13^C, ^15^N labeled, using different huPrP constructs and molar ratios between huPrP and Aβ(1-42) (**Table 1**).

### Characterization by density gradient ultracentrifugation, SDS-PAGE and RP-HPLC

For biophysical characterization of e. g. sample huPrP(23-144)-Aβ*, sucrose density gradient ultracentrifugation (DGC) was performed. To this end, 10 μl of the sedimented, but unwashed sample was diluted with 90 μl of 30 mM Tris-HCl buffer, pH 7.4 and applied on a discontinuous sucrose gradient (see ^33^) and centrifuged for 3 h at 259,000 x *g* and 4 °C. After fractionation, each of the 14 fractions was analyzed by Tris-Tricine SDS-PAGE and RP-HPLC as previously described^33^ (**Supplementary Fig. 3**).

RP-HPLC revealed the Aβ:huPrP(23-144) stoichiometry shown in **Table 1** as determined in a single measurement. Sample huPrP(23-144)_exc_-Aβ* was not separated by DGC but measured by RP-HPLC and revealed an Aβ:huPrP(23-144) stoichiometry of 3.7 ± 0.12 to 1 after fivefold measurement of the same sample. All stoichiometries represent monomer equivalents.

### Preparation of solution NMR samples

For the sequence-specific backbone resonance assignments samples of 0.36 mM uniformly 13C, ^15^N labeled huPrP(23-144) with 50 mM sodium acetate buffer in 10 % (v/v) D_2_O (pH 4.5) and 0.30 mM uniformly ^13^C, ^15^N labeled huPrP(23-144) with 50 mM HEPES buffer in 10 % (v/v) D_2_O (pH 7.0) were prepared as reported previously (Rösener et al.^33^). ^13^C-^13^C ‘TOtal Correlated SpectroscopY’ (TOCSY) NMR measurements in solution were performed on a sample containing 0.33 mM uniformly ^13^C, ^15^N labeled huPrP(23-144) monomers (Rösener et al.^33^) with 0.02 % (w/v) NaN_3_ in 30 mM HEPES buffer and 10 % (v/v) D_2_O (pH 6.7) and on a sample of 95 μM uniformly ^13^C, ^15^N labeled Aβ(1-42) (Isoloid GmbH) in 30 mM Tris-HCl buffer and 10 % (v/v) D_2_O (pH 7.2) at a temperature of 5.0 °C.

### Solid-state NMR experiments

The solid-state NMR measurements were performed either on Varian INOVA NMR spectrometers operating at field strengths of 14.1 Tesla (ω(^1^H)/(2π) = 600 MHz) for samples huPrP(23-144)*-Aβ, huPrP(23-230)*-Aβ and huPrP(23-144)-Aβ* or a Bruker AEON 18.8 Tesla (ω(^1^H)/(2π) = 800 MHz) spectrometer for sample huPrP(23-144)_exc_-Aβ*, equipped with 3.2 mm standard (Varian) or wide bore (Bruker) triple-resonance MAS probes. Therefore either 3.2 mm thick wall (25 μl, for samples huPrP(23-144)*-Aβ and huPrP(23-230)*-Aβ) or thin wall (36 μl, for sample huPrP(23-144)-Aβ*) rotors from Varian (Agilent) or 3.2 mm thick wall (46.7 μl, for sample huPrP(23-144)_exc_-Aβ*) rotors from Bruker were used. Sample temperatures were indirectly determined with an accuracy of ±5 °C for each spinning speed using nickelocene as an external reference^48^. Initial magnetization transfer from protons to ^13^C or ^15^N was either achieved by ‘insensitive nuclei enhanced by polarization transfer’ (INEPT)^49^ to selectively excite mobile regions via scalar coupling through-bond magnetization transfer from ^1^H to ^13^C (at ≈ 20, 27 or 30 °C) or by cross polarization (CP) (measured at ≈ 7, 0, −6 or −10 °C) via dipolar coupling through-space transfer for rigid parts. Additionally, several multi-dimensional homo- and heteronuclear correlation experiments for the assignment were recorded. Experimental details of all spectra recorded are given in **Supplementary Tables 2 to 6**. For homonuclear ^13^C-^13^C spectra, proton driven spin diffusion (PDSD)^50^ with mixing times between 10 to 300 ms was employed. Homonuclear double quantum correlation spectra were recorded with SPC5-recoupling^51^.

For site-specific assignment ^15^N-^13^C correlation spectra were recorded using SPECIFIC-CP^52^ for frequency selective polarization transfer from ^15^N to either ^13^Cα or ^13^CO and subsequent DARR-mixing. 2D NCA, NCACX and 3D NCACX and NCOCX spectra were used for the sequential walk through the backbone. During all acquisition and evolution times, high power broadband proton decoupling with SPINAL phase modulation^53^ (radio frequency intensity between 71 and 91 kHz) was used. All spectra were processed with NMRPipe^54^ with either squared and shifted sine bell or Gaussian window functions. The line width (FWHM) was estimated in 1D-slices (spectra processed with squared sine bell shifted by 0.35π or 0.40π) of 2D PDSD or NCACX/NCOCX spectra. ^13^C chemical shifts were externally referenced with adamantane by setting the low frequency signal of adamantane to 31.4 ppm on the DSS reference scale. ^15^N chemical shifts were indirectly referenced via the ^13^C chemical shifts. All resonances were assigned in CCPN ^55^. Integration of Aβ peaks was done in Topspin via the box sum method in a PDSD spectrum of huPrP(23-144)-Aβ*, measured at a temperature of ≈ 0 °C, a spinning frequency of 11 kHz and a mixing time of 50 ms.

### Solution NMR experiments

For the sequence-specific backbone resonance assignments of uniformly ^13^C, ^15^N labeled huPrP(23-144) in solution at pH 4.5, the following experiments were recorded at 5.0 °C on a Bruker AVANCE III HD 600 MHz NMR spectrometer equipped with an inverse triple resonance probe: 2D ^1^H-^15^N HSQC^56^, 3D HNCO^57^, and 3D HNCACB^58^ (Further experimental details are given in **Supplementary Table 7**). Sequence-specific backbone resonance assignments at pH 7.0 were obtained from 2D ^1^H-^15^N HSQC^56^, 3D HNCO^57^, 3D HNCACB^58^, and 3D BEST-TROSY-(H)N(COCA)NH^59^ experiments recorded at 5.0 °C on a Varian VNMRS 800 MHz NMR spectrometer equipped with an inverse triple resonance probe. A 2D ^13^C-^13^C TOCSY spectrum with a 13.6 ms 13.9 kHz FLOPSY-16 isotropic mixing scheme^60^ of 0.33 mM uniformly ^13^C, ^15^N labeled huPrP(23-144) at 5.0 °C was recorded on a Bruker AVANCE III HD 700 MHz NMR spectrometer equipped with an inverse triple resonance probe. Because of the comparatively low protein concentration, a 2D ^13^C-^13^C TOCSY spectrum with a 15.1 ms 15.6 kHz FLOPSY-16 isotropic mixing scheme^60^ of 95 μM uniformly ^13^C, ^15^N labeled Aβ(1-42) at 5.0 °C was recorded on a Bruker AVANCE III HD 800 MHz NMR spectrometer equipped with a ^13^C/^15^N observe triple resonance probe; a total of 1536 transients was collected over the course of three weeks and added up to further improve the signal-to-noise ratio. All triple resonance probes were cryogenically cooled and equipped with z axis pulsed field gradient capabilities. The sample temperature was calibrated using methanol-d_4_^61^. The ^1^H_2_O resonance was suppressed by gradient coherence selection with water flip-back^62^, with quadrature detection in the indirect dimensions achieved by States-TPPI^63^ and the echo-antiecho method^64,65^. All solution NMR spectra were processed with NMRPipe^54^ software and analyzed with NMRViewJ^66^ and CCPN^55^. ^1^H chemical shifts were referenced with respect to external DSS in D_2_O, ^13^C and ^15^N chemical shifts were referenced indirectly^67^. Random Coil Index (RCI)^40^ backbone order parameters, S_RCI_^2^, were calculated from the backbone chemical shifts using TALOS-N^68^ with the default parameters.

To obtain sequence-specific backbone resonance assignments for huPrP(23-144) at different pH values ranging from 4.5 to 7.0 and at a temperature of 5.0 °C, we employed the following strategy: (i) In the first step, as many resonance assignments as possible (see above) were transferred from huPrP(23-230) to the ^1^H-^15^N HSQC spectrum of huPrP(23-144) at pH 4.5 and 20.0 °C. (ii) Next, these resonance assignments were propagated along a temperature series of ^1^H-^15^N HSQC spectra of huPrP(23-144) at pH 4.5 recorded at temperatures of 15.0 °C, 10.0 °C, and 5.0 °C. (iii) The resulting sequence-specific backbone resonance assignments at pH 4.5 and 5.0 °C were verified and completed using HNCO and HNCACB triple-resonance experiments. (iv) These resonance assignments were then propagated along a pH series of ^1^H-^15^N HSQC spectra of huPrP(23-144) recorded at pH values of 5.3, 6.0, and 7.0 at a temperature of 5.0 °C. (v) Finally, the resulting sequence-specific backbone resonance assignments at pH 7.0 and 5.0 °C were verified and completed using HNCO, HNCACB, and BEST-TROSY-(H)N(COCA)NH triple-resonance experiments (**Supplementary Fig. 2**).

## Acknowledgments

The authors acknowledge access to the Jülich-Düsseldorf Biomolecular NMR Center that is jointly run by the Forschungszentrum Jülich and Heinrich-Heine-University Düsseldorf. The 800 MHz ^13^C/^15^N observe cryoprobe was funded by a grant from the German Research Foundation (DFG INST 208/620-1 FUGG) to D.W. L.G. and D.W. acknowledge support from the Russian Science Foundation (RSF) (project no. 20-64-46027). This work was supported by the Entrepreneur Foundation at the Heinrich-Heine-University of Düsseldorf and the DFG (HE 3243/4-1). Support from a European Research Council (ERC) Consolidator Grant (grant agreement no. 726368) to W.H. is acknowledged.

## Supplementary information

Supplementary information includes 23 Figures, and 7 tables.

## Data Availability

The assigned chemical shifts of huPrP(23-144) at pH 4.5 and pH 7.0 have been deposited with the Biological Magnetic Resonance Data Bank (BMRB) under accession codes 28115 and 28116, respectively.

## Author contributions

L.G and H.H. conceived the project and designed the experiments. A.S.K., N.S.R., P.N. and L.G. performed experiments, A.S.K., D.F., P.N. and H.H. analyzed data. A.S.K., D.W. and H.H. wrote the manuscript with input from all authors. All authors discussed the results and commented on the manuscript.

