## Supplementary information for "Structural details of amyloid beta oligomers in complex with human prion protein as revealed by solid-state MAS NMR spectroscopy"

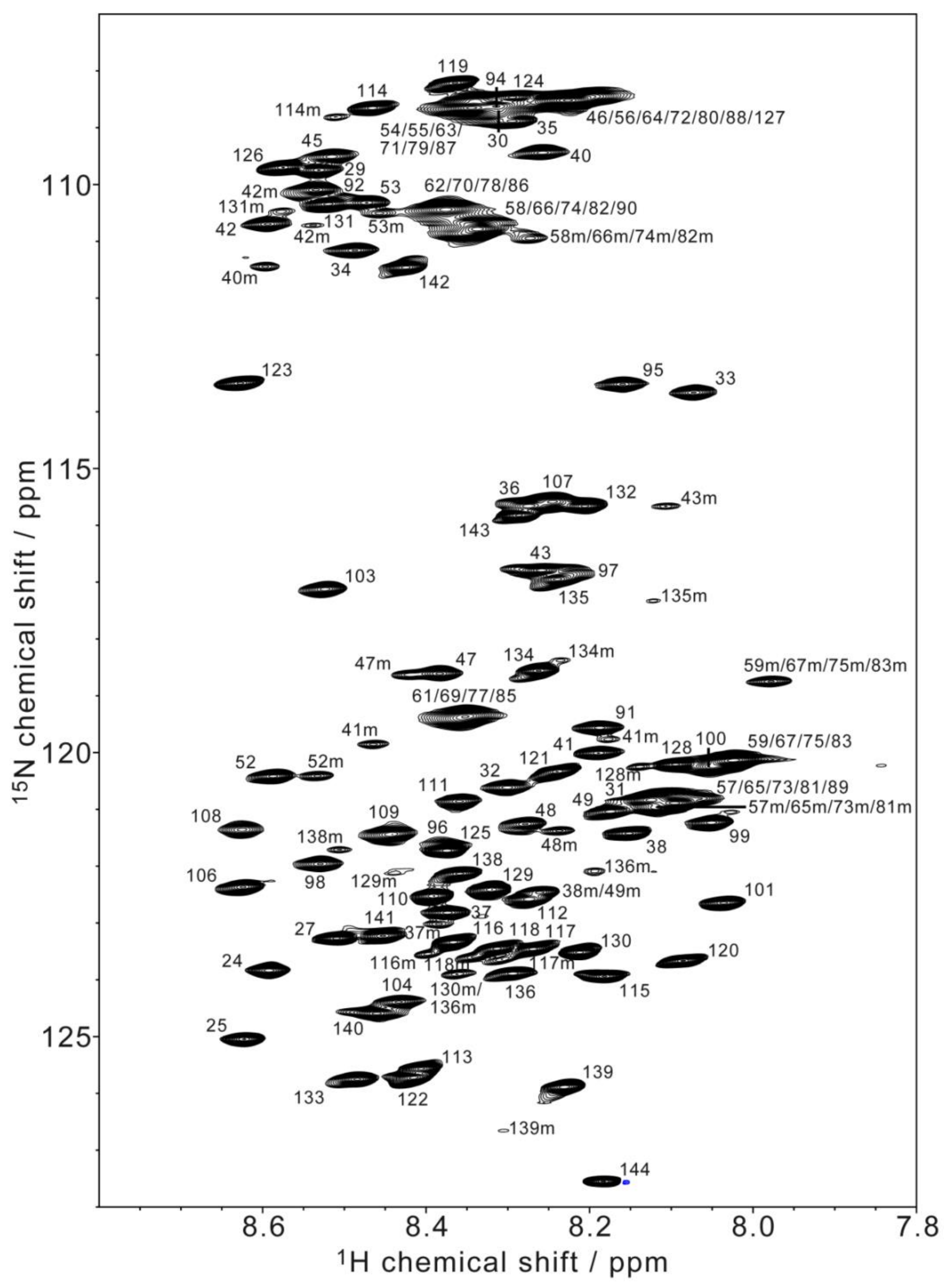

**Supplementary Fig. 1** Backbone amide region of the $^{1}H$-$^{15}N$ HSQC spectrum of a solution of 0.30 mM [U-$^{13}C$,$^{15}N$] huPrP(23-144) in 50 mM HEPES (pH 7.0), 10% (v/v) $D_2O$ recorded at 5.0 °C and 800 MHz (positive contours black, negative contours blue). Backbone resonance assignments are indicated by residue numbers, minor resonances (e. g. due to proline residues in *cis* conformation or methionine oxidation) are indicated by an m. Four octarepeats (P51 to Q91) share the same sequence (**Fig. 2a**) and could not be resolved. The backbone amide resonance of G93 could not be identified due to severe resonance overlap.

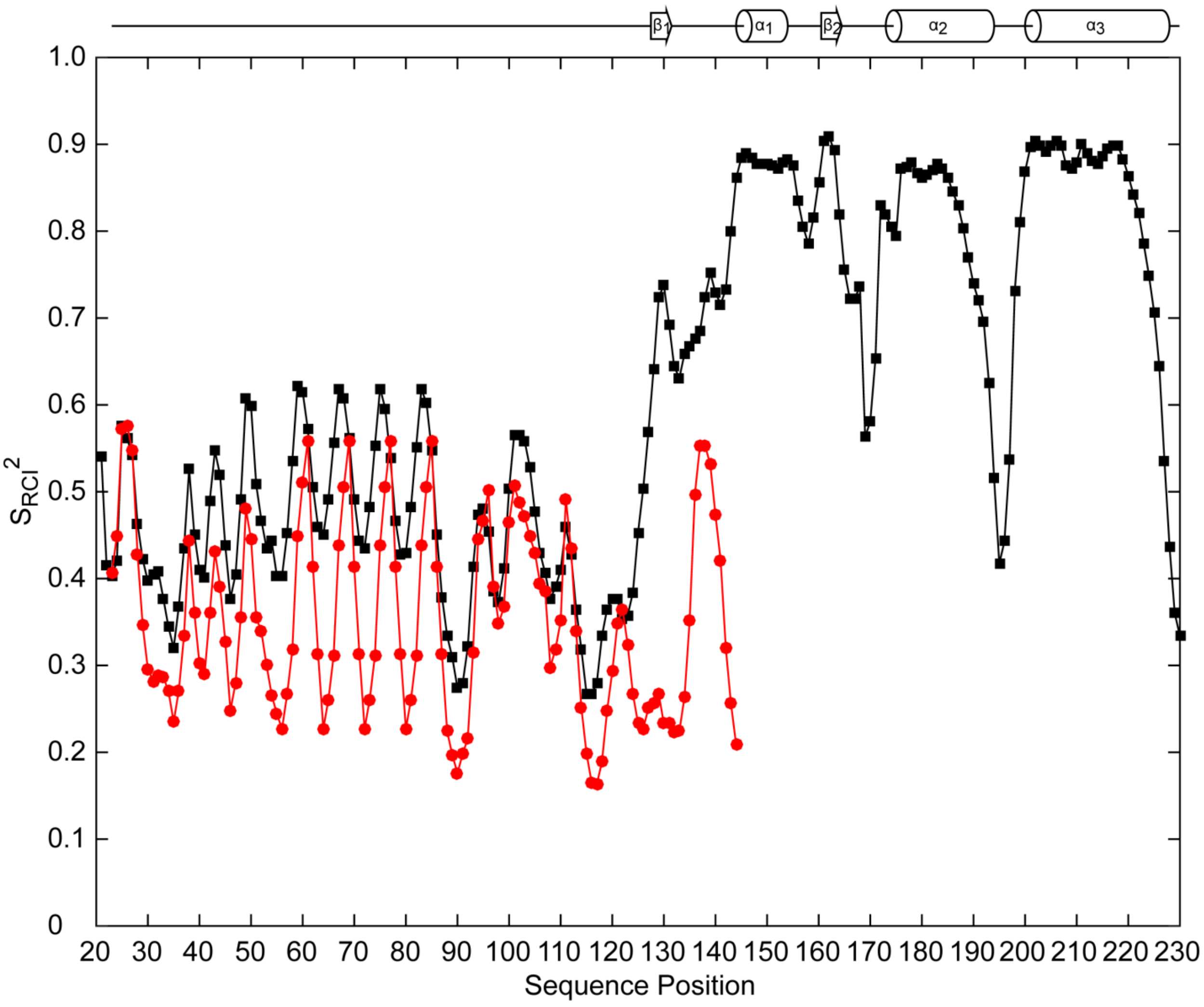

**Supplementary Fig. 2** Backbone order parameters predicted from the Random Coil Index (RCI), $S_{RCI}^2$, for huPrP(23-230) at pH 4.5 (BMRB 4402, ref. [1], black) and huPrP(23-144) at pH 7.0 (BMRB 28116, this work, red), as calculated by TALOS-N[2] with the default parameters. Assigned chemical shifts for the short N-terminal cloning artifact (Gly for huPrP(23-144), Gly-Ser for huPrP(23-230)) were included. Full-length huPrP(23-230) consists of a highly disordered N-terminal region comprising residues 23 to 124 with chemical shifts very close to random coil values and concomitantly low $S_{RCI}^2$ values below about 0.6 (black), and a globular C-terminal prion domain comprising residues 125 to 228, whose regular secondary structure elements are indicated above the figure. Upon truncation, residues 125 to 144 of huPrP(23-144) also become disordered, with chemical shifts very close to random coil values (**Supplementary Fig. 1**) and concomitantly low $S_{RCI}^2$ values below ≈ 0.6 for all residues from 23 to 144 (red).

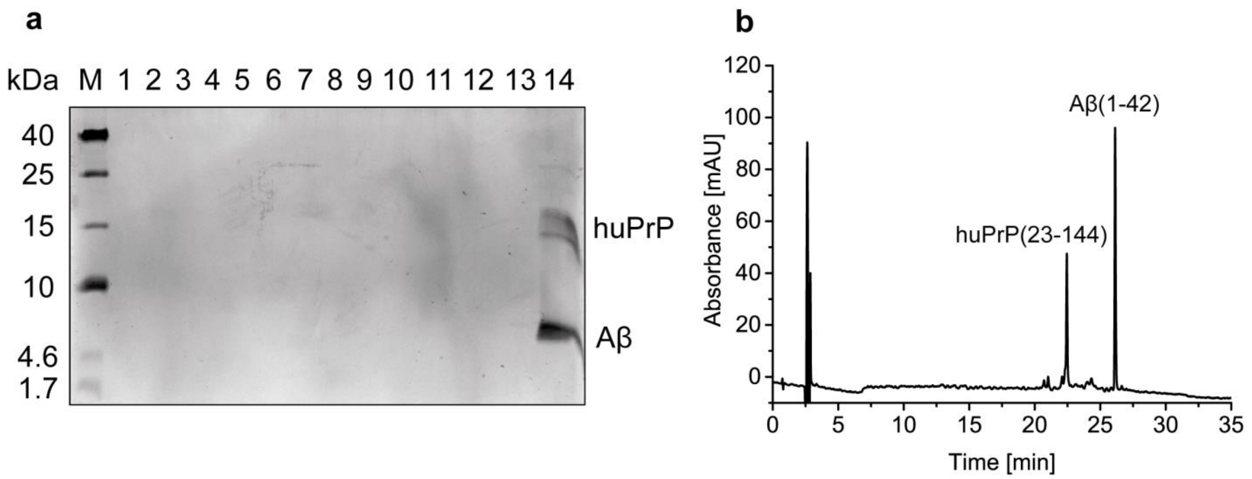

**Supplementary Fig. 3 a** Sucrose DGC[3] of a tenfold dilution of huPrP(23-144)-Aβ* (* species is $^{13}$C, $^{15}$N uniformly labeled) before washing shows huPrP and Aβ only in fraction 14, corresponding to the highest density[4]. **b** Quantitative analysis by RP-HPLC on fraction 14 of the DGC revealed an Aβ(1-42) to huPrP(23-144) stoichiometry of 8.6 to 1 (monomer equivalents, single measurement).

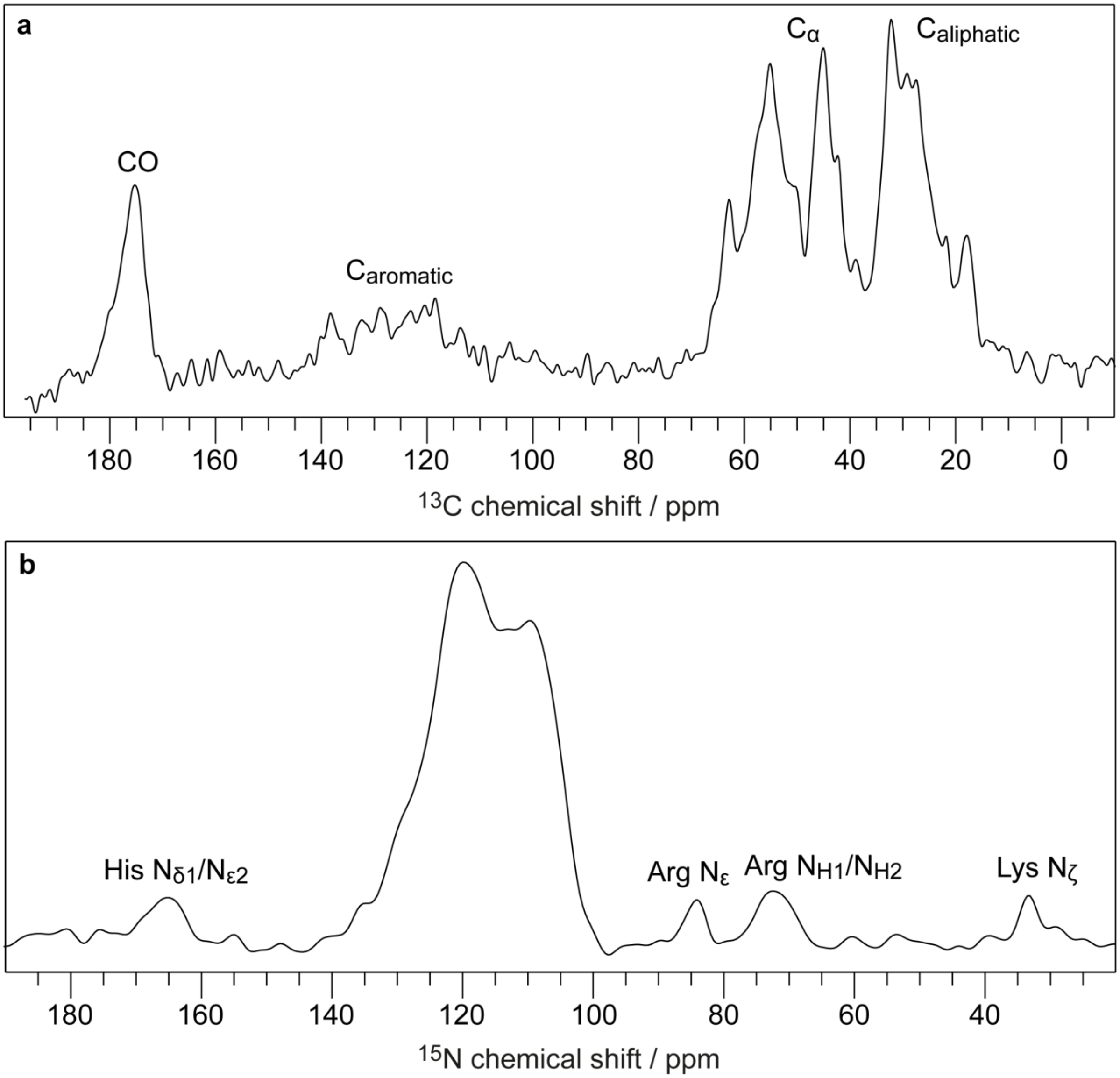

**Supplementary Fig. 4 a** $^{1}$H-$^{13}$C CP spectrum of huPrP(23-144)*-Aβ (* species is $^{13}$C, $^{15}$N uniformly labeled), measured at a temperature of ≈ 0 °C, at a spinning frequency of 11 kHz and 256 scans. **b** $^{1}$H-$^{15}$N CP spectrum of huPrP(23-144)*-Aβ, measured at a temperature of ≈ -6 °C, a spinning frequency of 11 kHz and 2000 scans.

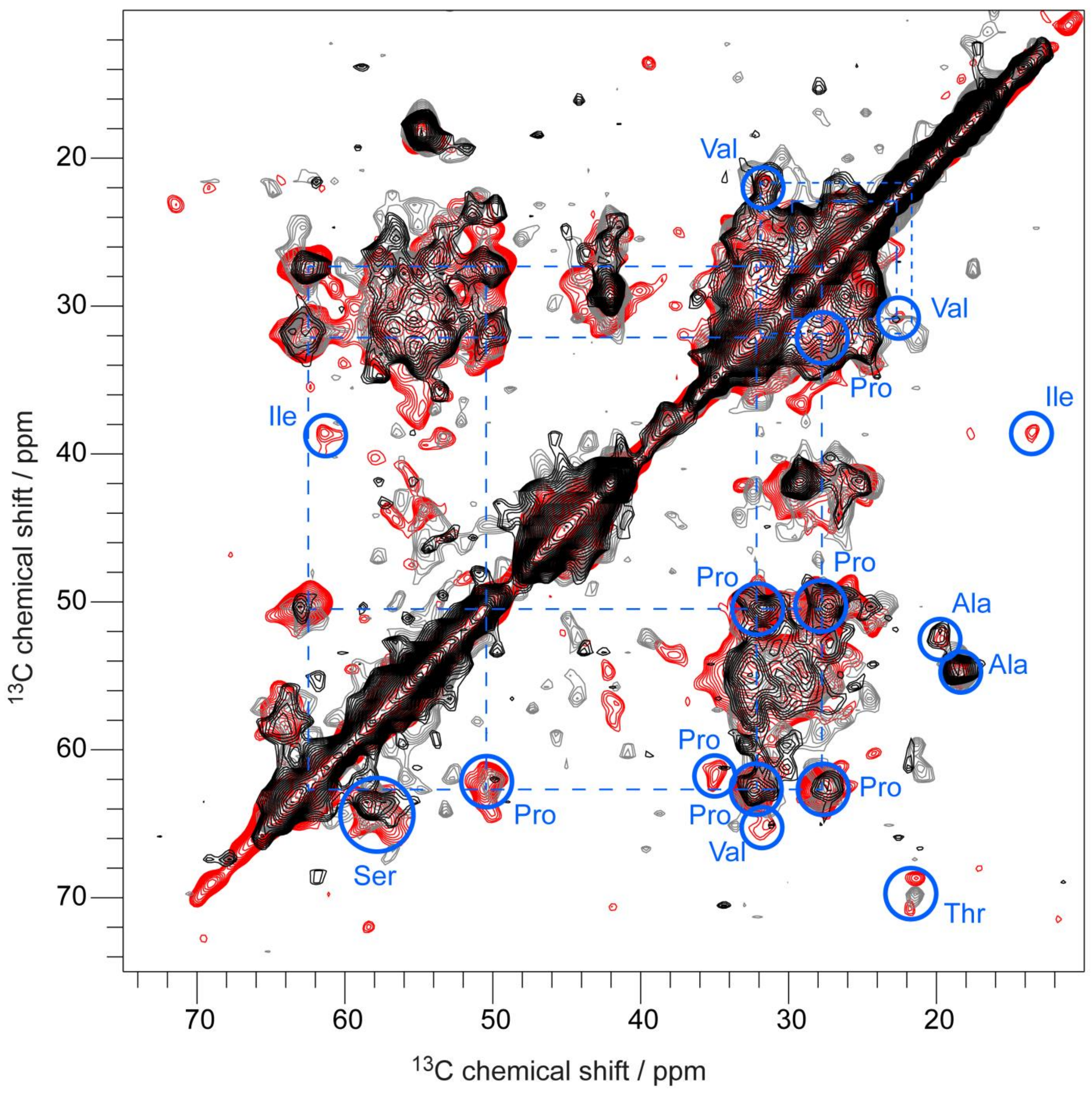

**Supplementary Fig. 5** PDSD spectra of huPrP(23-144)*-Aβ (* species is $^{13}$C, $^{15}$N uniformly labeled), measured at a spinning frequency of 11 kHz, a temperature of ≈ -6 °C and a mixing time of 30 ms (red) or a temperature of ≈ 0 °C and a mixing time of 50 ms (black) or 100 ms (grey). Blue circles indicate some identified amino acid types, dashed lines Pro and Val connections.

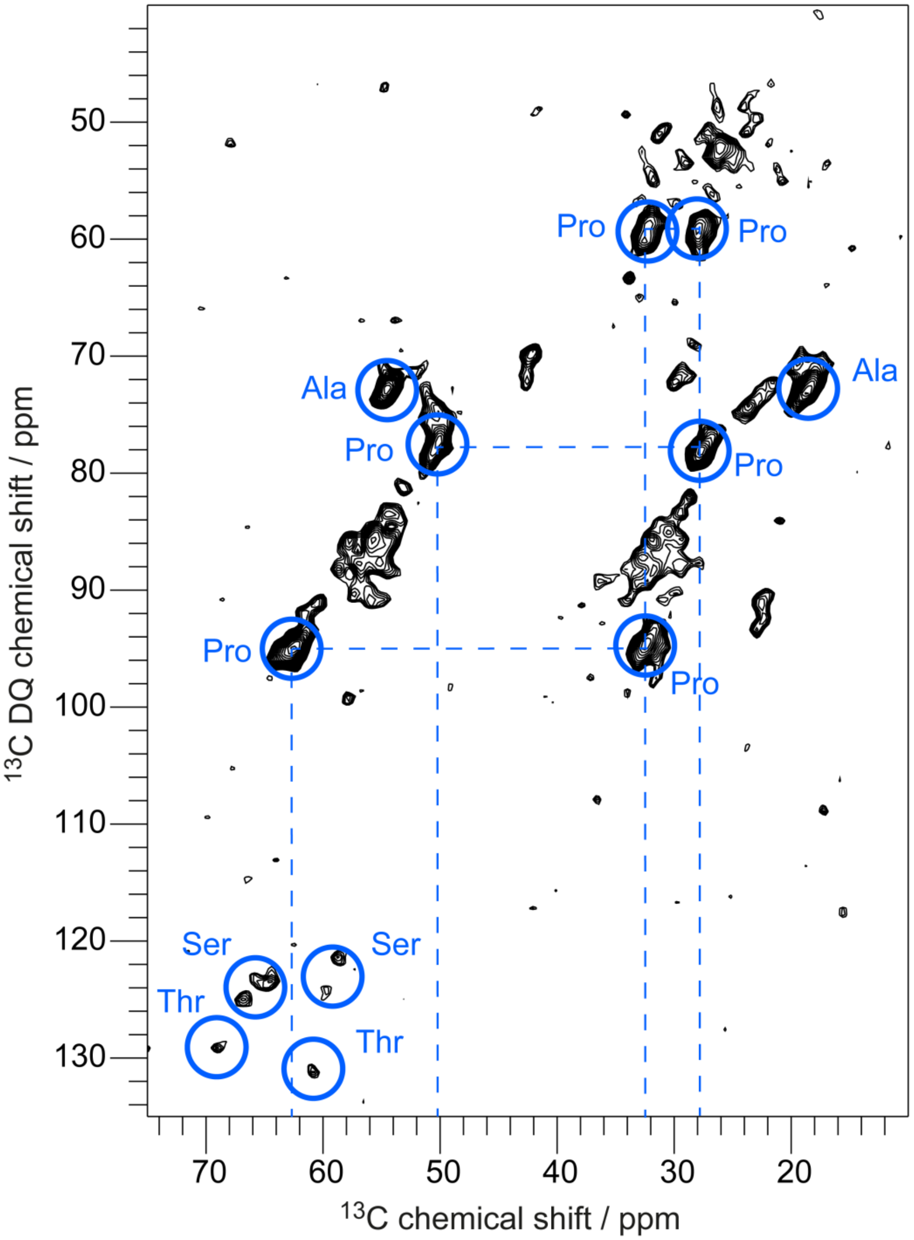

**Supplementary Fig. 6** Double-quantum (DQ) correlation spectrum of huPrP(23-144)*-Aβ (* species is $^{13}$C, $^{15}$N uniformly labeled) with SPC5-recoupling, measured at a temperature of ≈ -6 °C and a spinning frequency of 8 kHz. Blue circles indicate identified amino acid types, dashed lines Pro connections.

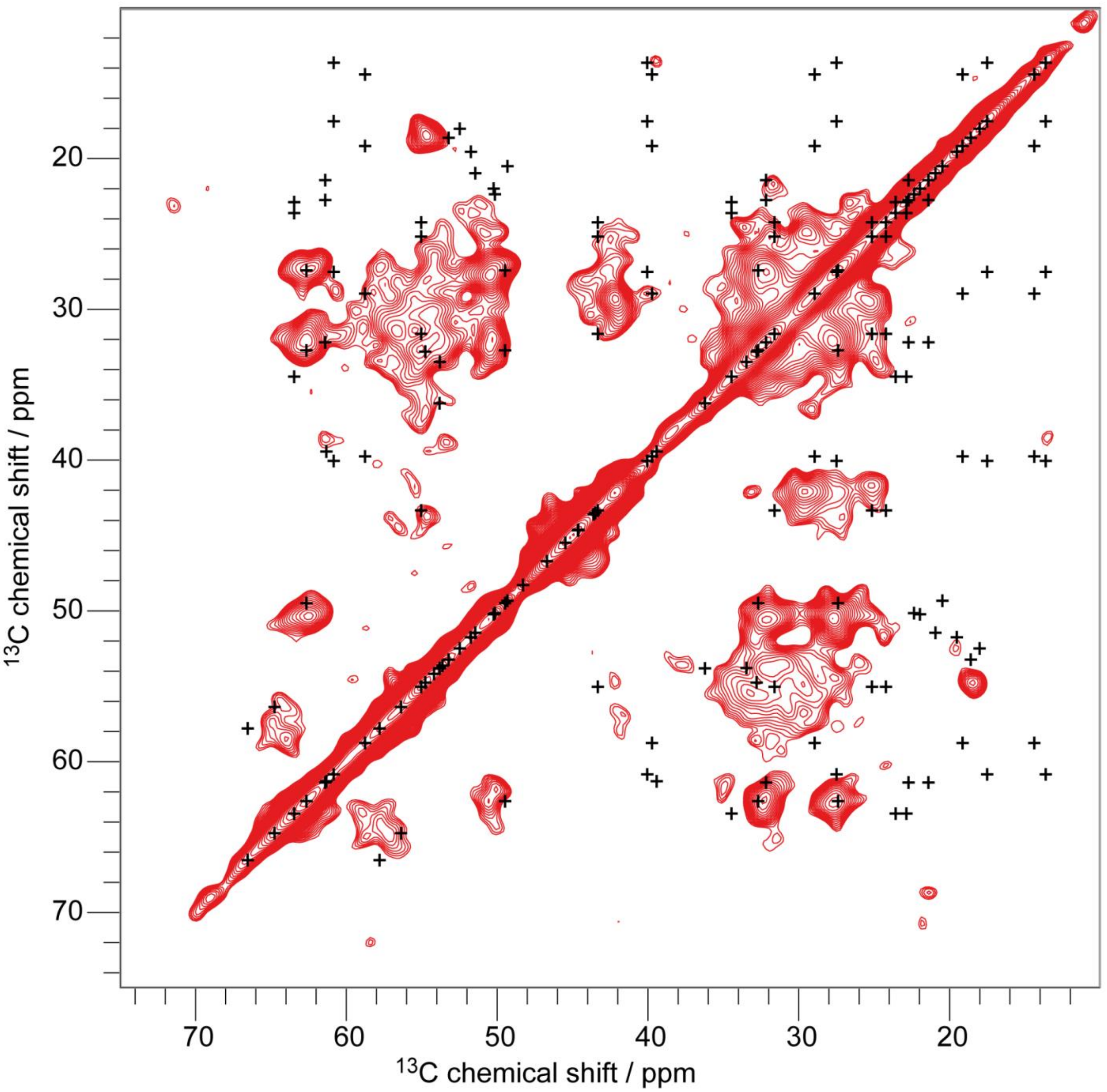

**Supplementary Fig. 7** Comparison of a PDSD spectrum (recorded at a temperature of ≈ -6 °C, a spinning frequency of 11 kHz and a mixing time of 30 ms, same as in **Fig.1**) of huPrP(23-144)*-Aβ (* species is $^{13}$C, $^{15}$N uniformly labeled) with the chemical shifts of a huPrP(23-144) fibril measured by Theint et al. (BMRB entry 26925)[5], shown as black crosses.

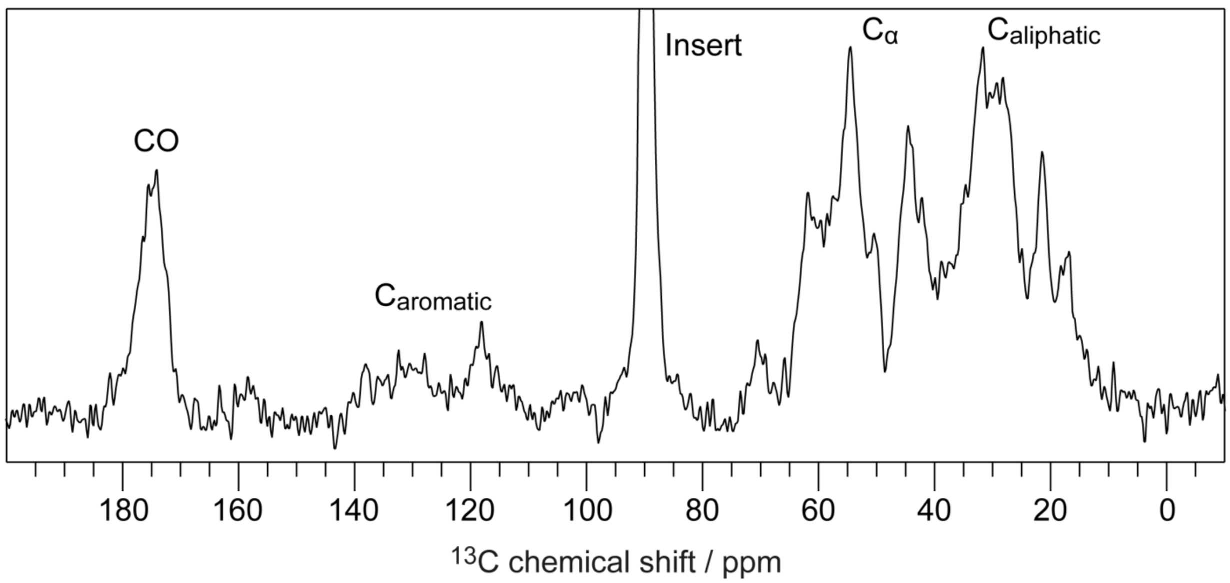

**Supplementary Fig. 8** $^{1}H$-$^{13}C$ CP spectrum of huPrP(23-230)*-Aβ (* species is $^{13}C$, $^{15}N$ uniformly labeled), measured at a temperature of ≈ -10 °C, a spinning frequency of 11 kHz and 512 scans. The signal at 90 ppm belongs to a break-proof rotor insert and is cut off for clarity. The insert was used as a precaution because at the beginning of the measurements it was not known if PrP in huPrP(23-230)*-Aβ was present in its pathogenic $PrP^{Sc}$ conformation.

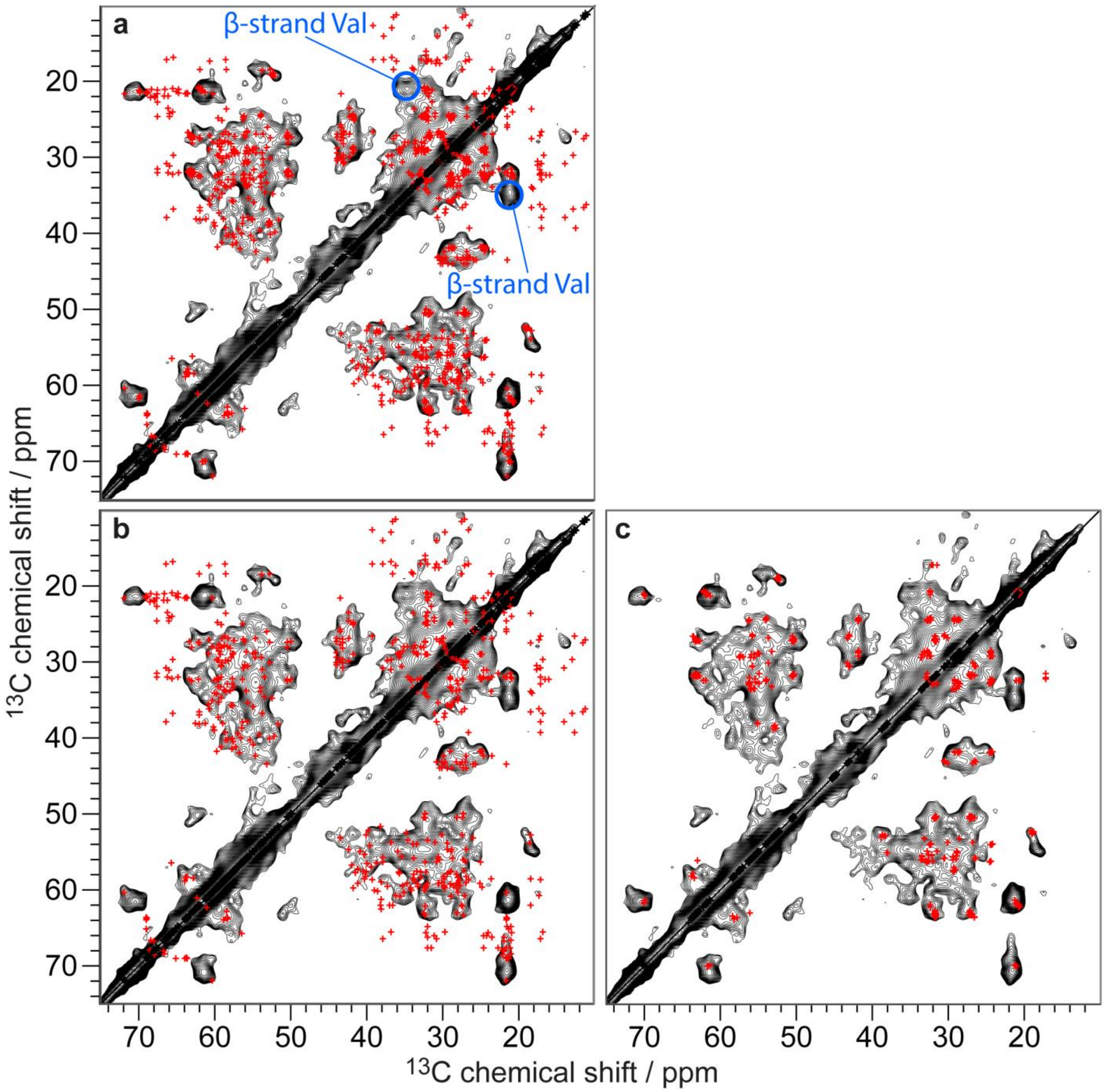

**Supplementary Fig. 9** Comparison of a PDSD spectrum of huPrP(23-230)*-Aβ (black) (* species is $^{13}C$, $^{15}N$ uniformly labeled) to predicted peaks, indicated as red crosses. Measured chemical shifts of Zahn et al. (BMRB entry 4402) were used for the prediction. For those carbon atoms in the structured C terminus (starting from residue 125) for which no chemical shift assignments were made by Zahn et al., chemical shifts were predicted from the PDB-structure of monomeric huPrP (PDB-Entry 1QLZ)[1] with SHIFTX2[6]. For residues from the unstructured N-terminal part for which no chemical shift assignments were made by Zahn et al., random coil BMRB values[7] were used. Cross peaks of up to two bonds were included in the prediction. Full spin-systems were simulated, but at 30 ms mixing time not all cross-correlations necessarily show up with sufficient intensity in the experimental spectrum. **a** Values for residues MK23-

S230, **b** L125-S230 (C terminus only), and **c** MK23-G124 (N terminus only) were used. The separation between N-terminal and C-terminal regions is made between G124 and L125, where the ordered region of the solution NMR structure starts (**Fig. 2a**). The PDSD spectrum was measured at a temperature of ≈ 0 °C, a spinning frequency of 11 kHz and a mixing time of 30 ms. β-strand-like Val is indicated with blue circles, it is the only peak in the spectrum of huPrP(23-230)*-Aβ which is not superimposed at all. Note that these structural changes are not due to the different pH of huPrP(23-230)*-Aβ (pH 7.4) and soluble huPrP(23-230) (pH 4.5), as the conformation of the globular domain (residues 125 to 230) of monomeric huPrP(23-230) at pH 4.5 and pH 7.0 is extremely similar[8].

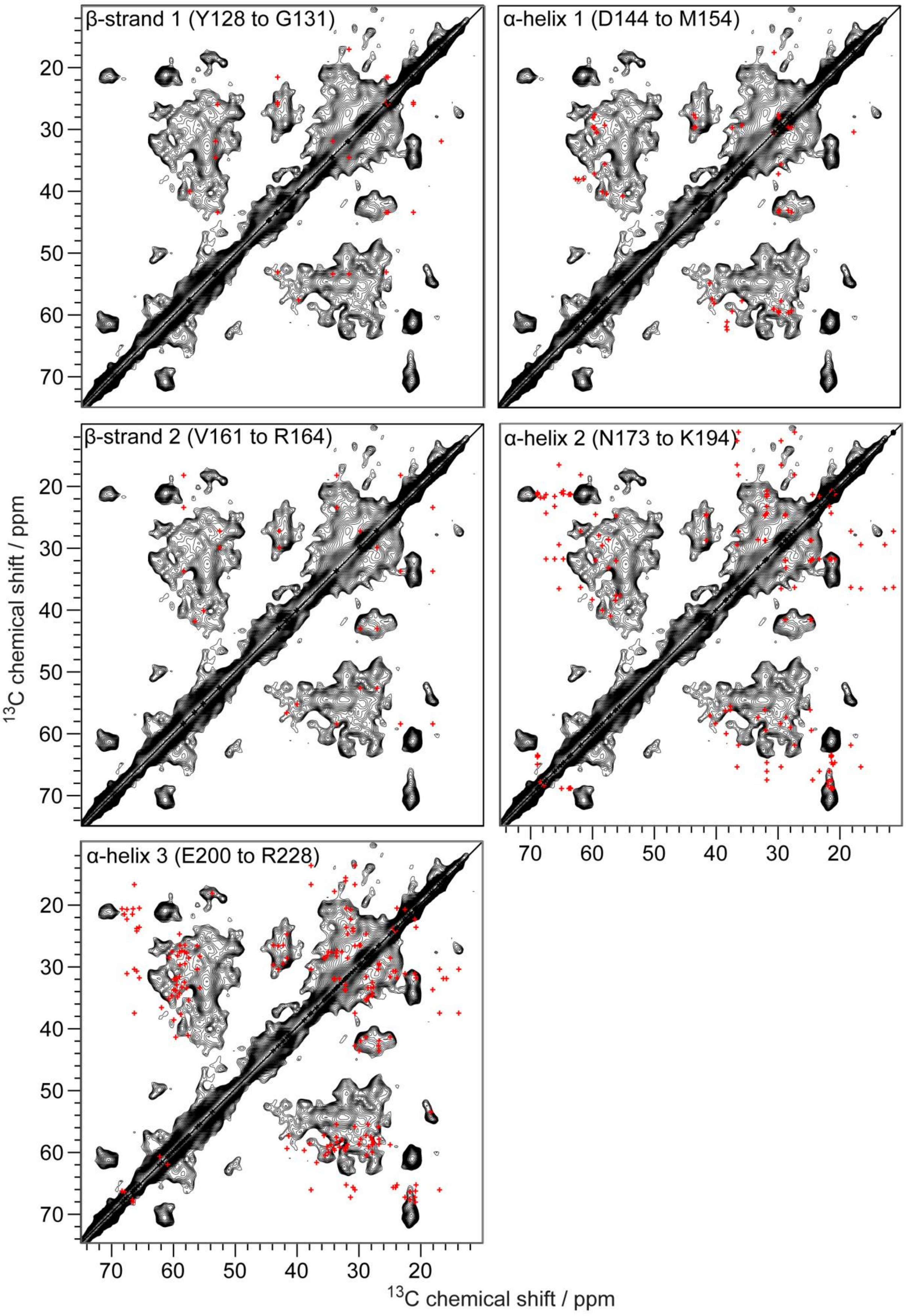
β-strand 1 (Y128 to G131)
α-helix 1 (D144 to M154)
β-strand 2 (V161 to R164)
α-helix 2 (N173 to K194)
α-helix 3 (E200 to R228)
13C chemical shift / ppm
13C chemical shift / ppm

**Supplementary Fig. 10** Comparison of a PDSD spectrum of huPrP(23-230)*-Aβ (black) (* species is $^{13}C$, $^{15}N$ uniformly labeled) to predicted peaks, indicated as red crosses. Measured chemical shifts of Zahn et al. (BMRB entry 4402) were used for the prediction. For those carbon atoms in the structured C terminus (starting from residue 125) for which no chemical shift assignments were made by Zahn et al., chemical shifts were predicted from the PDB-structure of monomeric huPrP(23-230) (PDB-Entry 1QLZ)[1] with SHIFTX2[6]. Cross peaks of up to two bonds were included in the prediction. Full spin-systems were simulated, but at 30 ms mixing time not all cross-correlations necessarily show up with sufficient intensity in the experimental spectrum. (β-strand 1) Values for residues Y128 to G131, (α-helix 1) D144 to M154, (β-strand 2) V161 to R164, (α-helix 2) N173 to K194 and (α-helix 3) E200 to R228 were used. The PDSD spectrum was measured at a temperature of ≈ 0 °C, a spinning frequency of 11 kHz and a mixing time of 30 ms. Except for longer correlations over three or four bonds, some side chains of Leu, Met and V161, the resonances of the two β-strands and the first α-helix determined by Zahn et al. align well with the resonances of huPrP(23-230)*-Aβ. But as these resonances determined by Zahn et al. overlap with other resonances in the spectrum of huPrP(23-230)*-Aβ, it is possible that they are either not visible or shifted. Therefore, no conclusion can be drawn about the conservation of the two β-strands and the first α-helix. For the last two α-helices also longer correlations are missing. But interestingly all α-helical like Ile, Val and, in the second α-helix, Thr resonances are missing, too. This means Ile, Val and Thr are more random coil or β-strand like in huPrP(23-230)*-Aβ or partly undetectable, i.e. flexible on an intermediate time scale (as no signals are observed in an INEPT experiment (data not shown)).

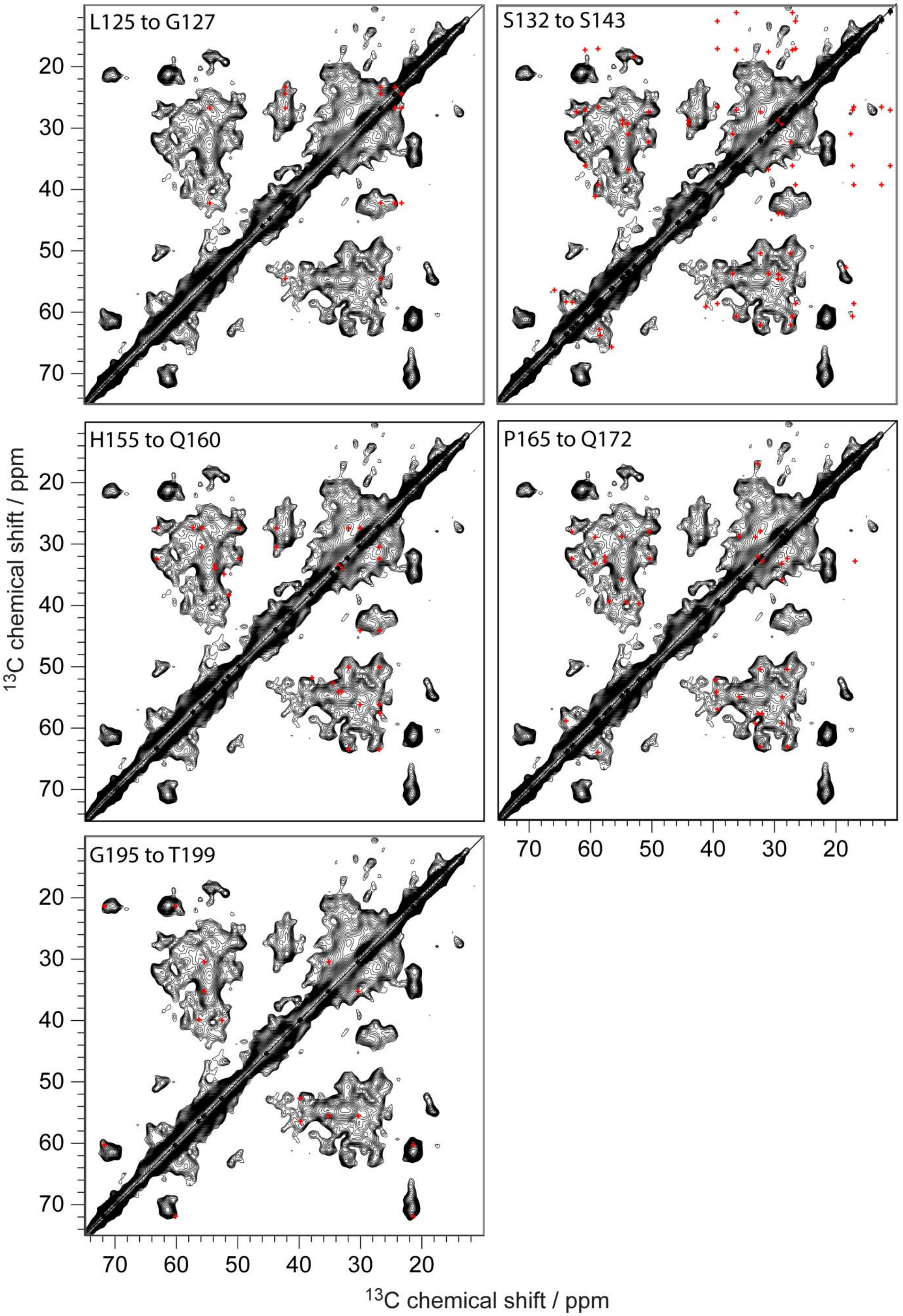
L125 to G127
S132 to S143
H155 to Q160
P165 to Q172
G195 to T199
20
30
40
50
60
70
$^{13}$C chemical shift / ppm
70
60
50
40
30
20
$^{13}$C chemical shift / ppm

**Supplementary Fig. 11** Comparison of a PDSD spectrum of huPrP(23-230)*-Aβ (black) (* species is $^{13}C$, $^{15}N$ uniformly labeled) to predicted peaks, indicated as red crosses. Measured chemical shifts of Zahn et al. (BMRB entry 4402) were used for the prediction. For those carbon atoms in the structured C terminus (starting from residue 125) for which no chemical shift assignments were made by Zahn et al., chemical shifts were predicted from the PDB-structure of monomeric huPrP (PDB-Entry 1QLZ)[1] with SHIFTX2[6]. Cross peaks of up to two bonds were included in the prediction. Full spin-systems were simulated, but at 30 ms mixing time not all cross-correlations necessarily show up with sufficient intensity in the experimental spectrum. Here it is shown for all loop regions between the β-strands and α-helices, namely L125 to G127, S132 to S143, H155 to Q160, P165 to Q172 and G195 to T199. The PDSD spectrum was measured at a temperature of ≈ 0 °C, a spinning frequency of 11 kHz and a mixing time of 30 ms. In the loop regions between the β-strands and α-helices only correlations over two or more bonds plus S143 are missing or shifted.

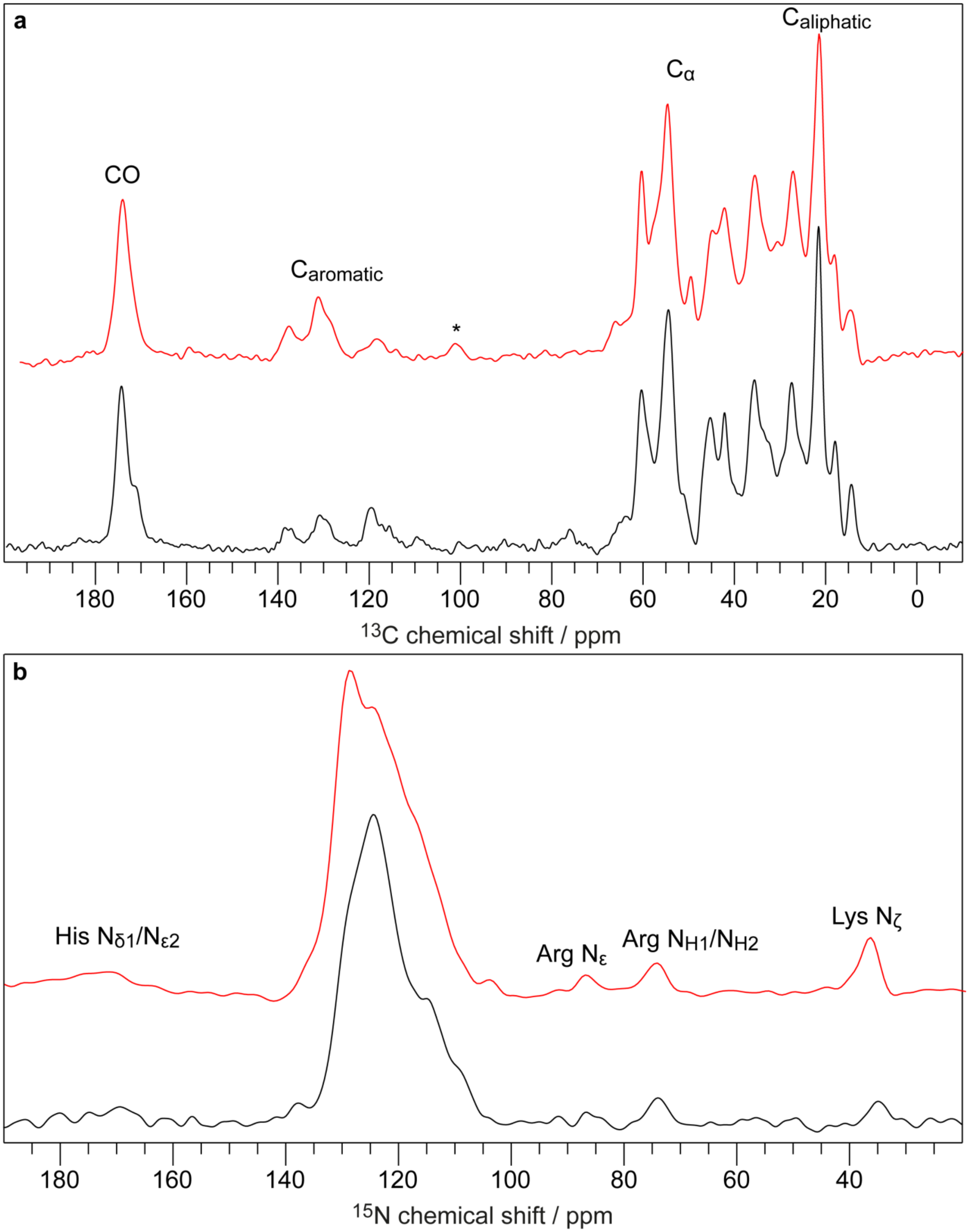

**Supplementary Fig. 12 a** (red) $^{1}$H-$^{13}$C CP spectrum of huPrP(23-144)-Aβ* (* species is $^{13}$C, $^{15}$N uniformly labeled), measured at a temperature of ≈ 0 °C, at a spinning frequency of 11 kHz and 128 scans. Spinning side bands are marked with an asterisk. **a** (black) $^{1}$H-$^{13}$C CP spectrum of huPrP(23-144)$_{exc}$-Aβ*

($_{exc}$ huPrP is in excess; * species is $^{13}C$, $^{15}N$ uniformly labeled), measured at a temperature of ≈ 7 °C, at a spinning frequency of 5 kHz and 256 scans. **b** (red) $^{1}H$-$^{15}N$ CP spectrum of huPrP(23-144)-Aβ*, measured at a temperature of ≈ 0 °C, a spinning frequency of 11 kHz and 2000 scans. **b** (black) $^{1}H$-$^{15}N$ CP spectrum of huPrP(23-144)$_{exc}$-Aβ*, measured at a temperature of ≈ 0 °C, a spinning frequency of 11 kHz and 2048 scans.

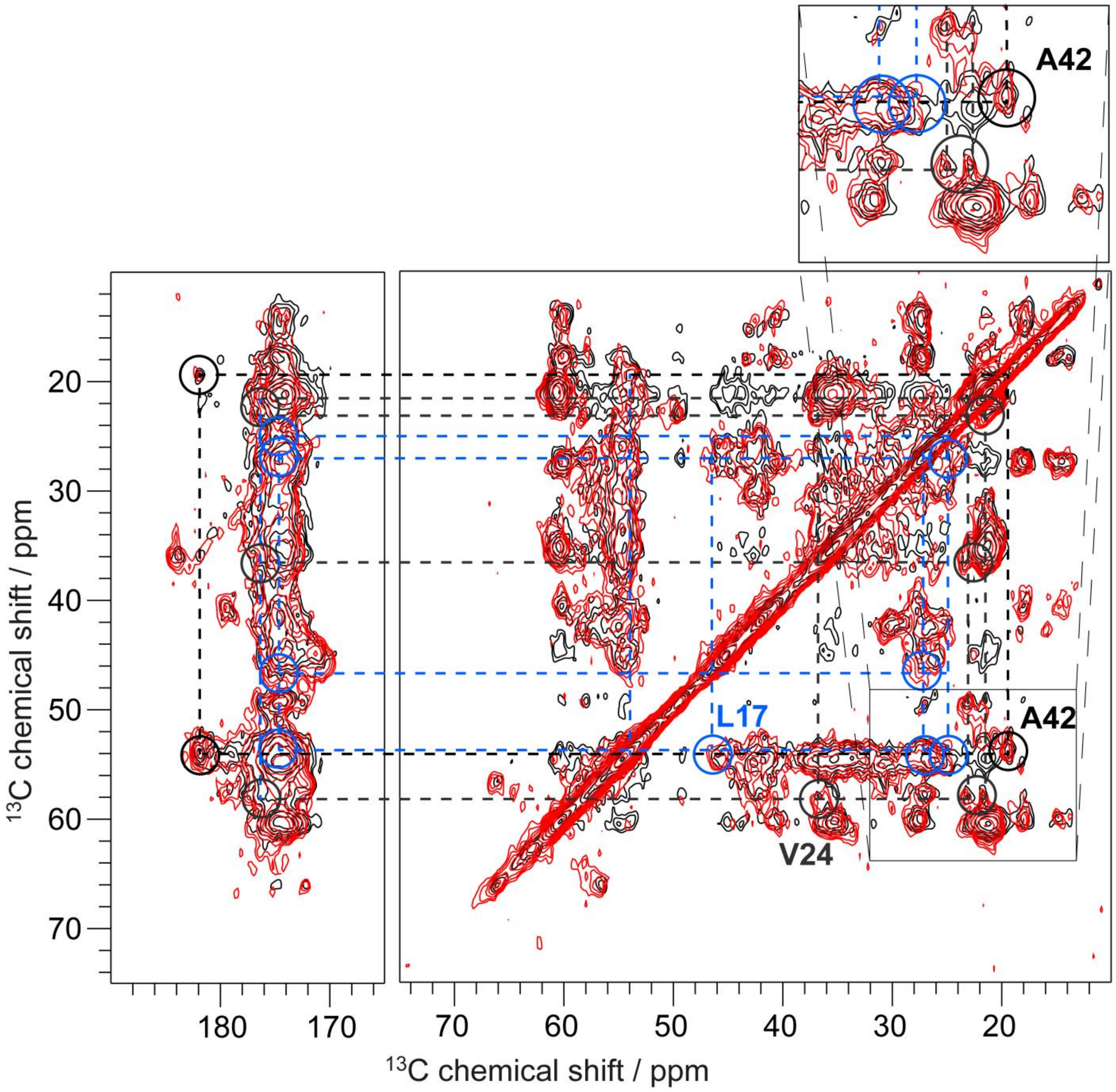

**Supplementary Fig. 13** Two PDSD spectra of huPrP(23-144)-Aβ* (* species is $^{13}$C, $^{15}$N uniformly labeled), measured at a temperature of ≈ 0 °C, a spinning frequency of 11 kHz and a mixing time of 50 ms (red) and 200 ms (black). Three identified residues are shown exemplarily with colored circles and dashed lines: L17 (blue), V24 (grey) and A42 (black). For L17 $C_{\delta1}$ and $C_{\delta2}$ chemical shifts are not separated due to the low signal dispersion. In contrast, V24 shows two separated $C_{\gamma1}$ and $C_{\gamma2}$ chemical shifts, which has been also observed in the Aβ(1-42) fibril polymorph of Gremer et al.[9]. Note the high CO chemical shift of A42 (182 ppm) (left part, black circles). This can be explained by a free and deprotonated state of the C-terminal carboxyl group. At 200 ms mixing time several multi-bond and inter-residual correlations appear, which are not visible at 50 ms mixing time, for example for residues Q15 to F20. As these residues additionally do not show multiple peaks, this is the structurally most conserved part in the oligomer. Altogether, these characteristics made it possible to assign these resonances to the appropriate residue in the amino acid sequence.

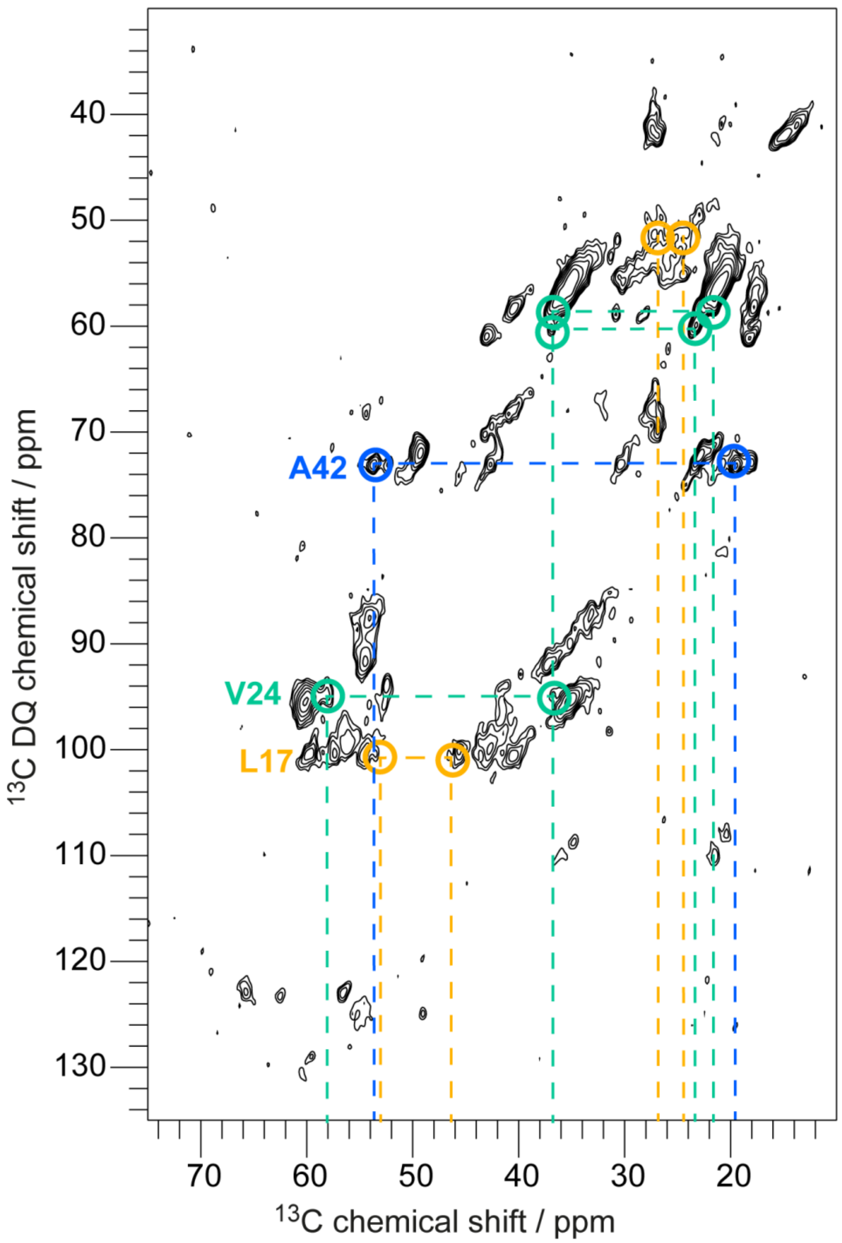

**Supplementary Fig. 14** Double-quantum (DQ) correlation spectrum of huPrP(23-144)-Aβ* (* species is $^{13}C$, $^{15}N$ uniformly labeled) with SPC5-recoupling, measured at a temperature of ≈ 0 °C and a spinning frequency of 8 kHz. Three identified residues are shown exemplarily with colored circles and dashed lines: L17 (orange), V24 (green) and A42 (blue).

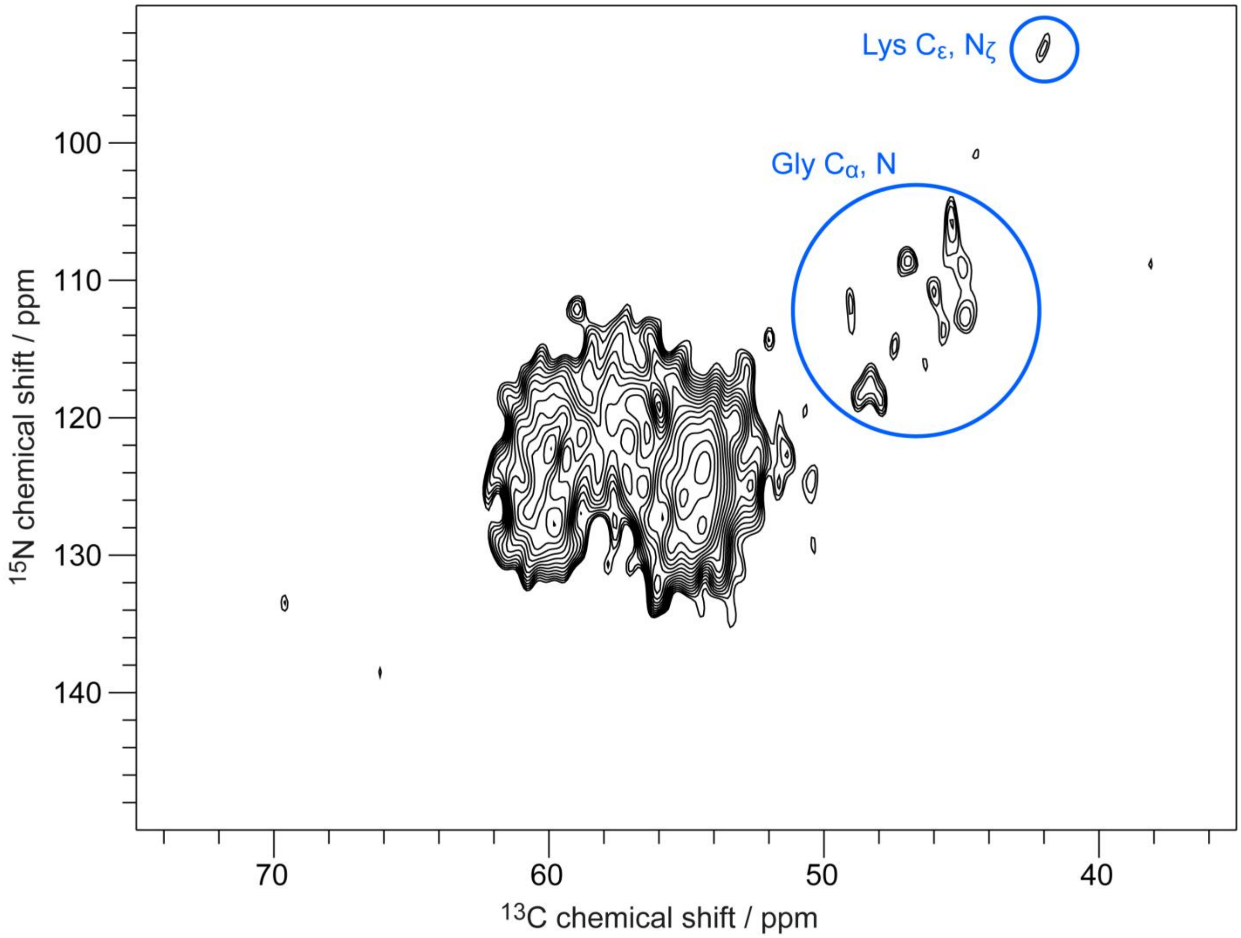

**Supplementary Fig. 15** NCA spectrum of huPrP(23-144)-Aβ* (* species is $^{13}$C, $^{15}$N uniformly labeled), measured at a temperature of ≈ 0 °C and a spinning frequency of 11 kHz. Blue circles indicate some identified amino acids.

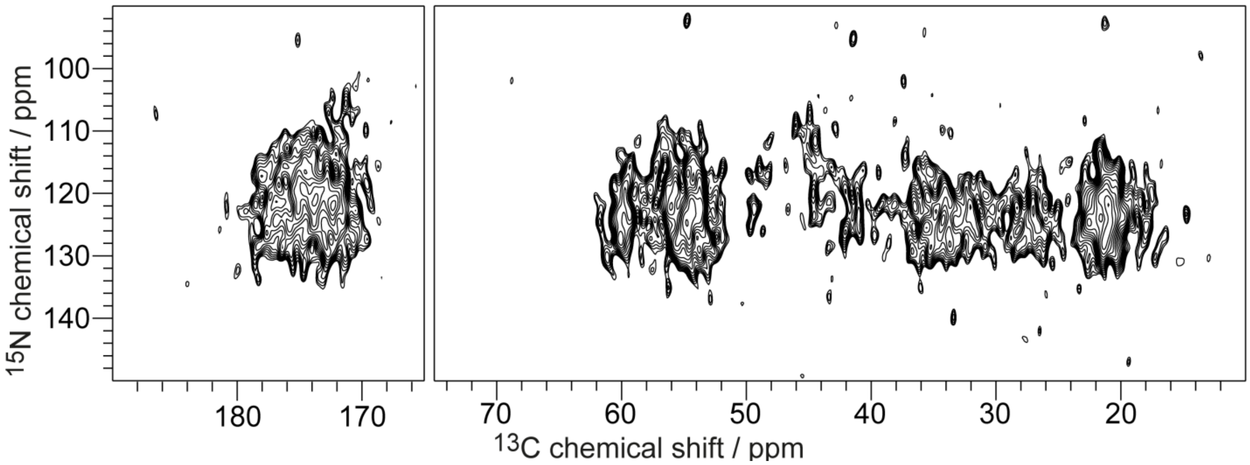

**Supplementary Fig. 16** 2D NCACX spectrum of huPrP(23-144)-Aβ* (* species is $^{13}$C, $^{15}$N uniformly labeled), measured at a temperature of ≈ 0 °C and a spinning frequency of 11 kHz.

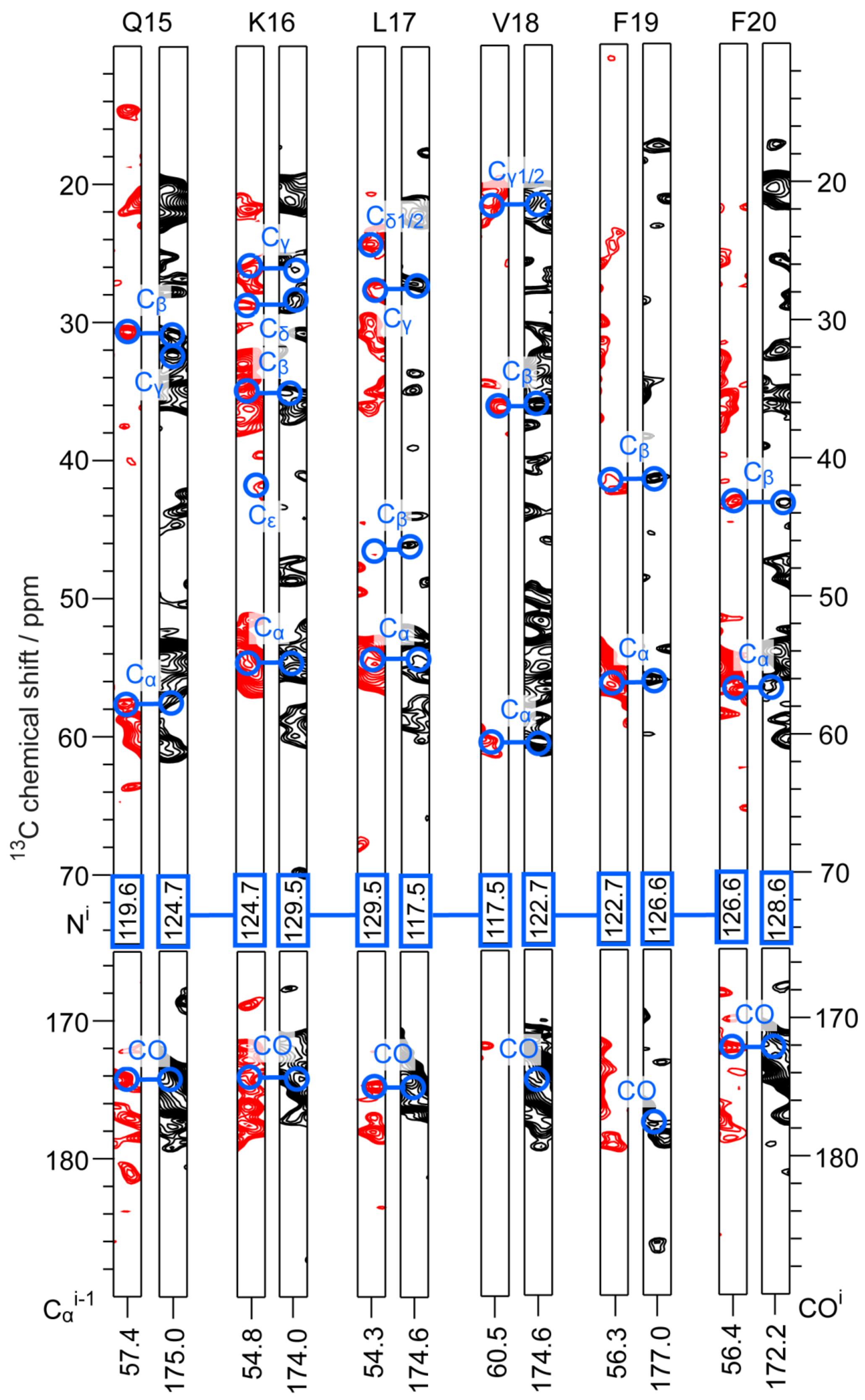

Q15
K16
L17
V18
F19
F20
13C chemical shift / ppm
Cα
Cβ
Cγ
Cδ
Cε
Cγ1/2
Cδ1/2
CO
Ni
119.6 124.7 124.7 129.5 129.5 117.5 117.5 122.7 122.7 126.6 126.6 128.6
Cαi-1
COi
57.4 175.0 54.8 174.0 54.3 174.6 60.5 174.6 56.3 177.0 56.4 172.2

**Supplementary Fig. 17** Representative strip plot of huPrP(23-144)-Aβ* (* species is $^{13}C$, $^{15}N$ uniformly labeled), 2D slices of 3D-NCACX (red) and 3D-NCOCX (black) spectra, measured at a temperature of ≈ 0 °C, a spinning frequency of 11 kHz and subsequent DARR-mixing with 60 (NCACX) or 70 (NCOCX) ms mixing time. The sequential walk for residues Q15 to F20 is shown in blue.

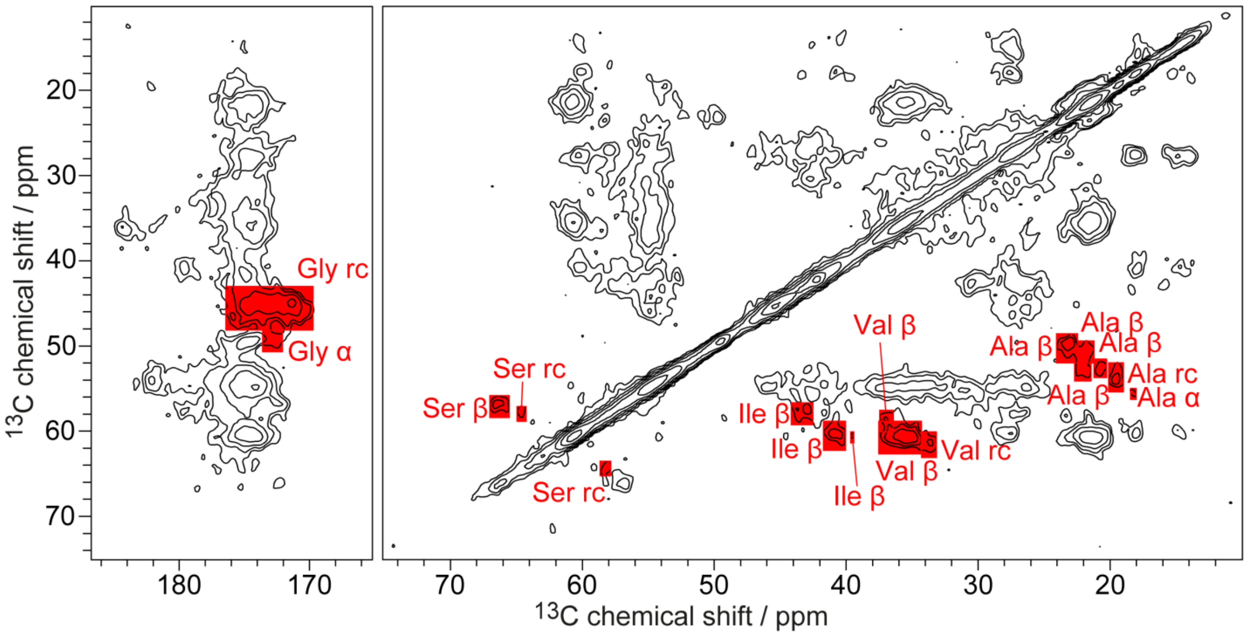

**Supplementary Fig. 18** 2D PDSD spectrum of huPrP(23-144)-Aβ* (* species is $^{13}C$, $^{15}N$ uniformly labeled), measured at a temperature of ≈ 0 °C, a spinning frequency of 11 kHz and a mixing time of 50 ms. α-helical (labeled with 'α'), unstructured (labeled with 'rc', random coil) and β-strand like (labeled with 'β') conformations were integrated by the box sum method in Topspin (red boxes).

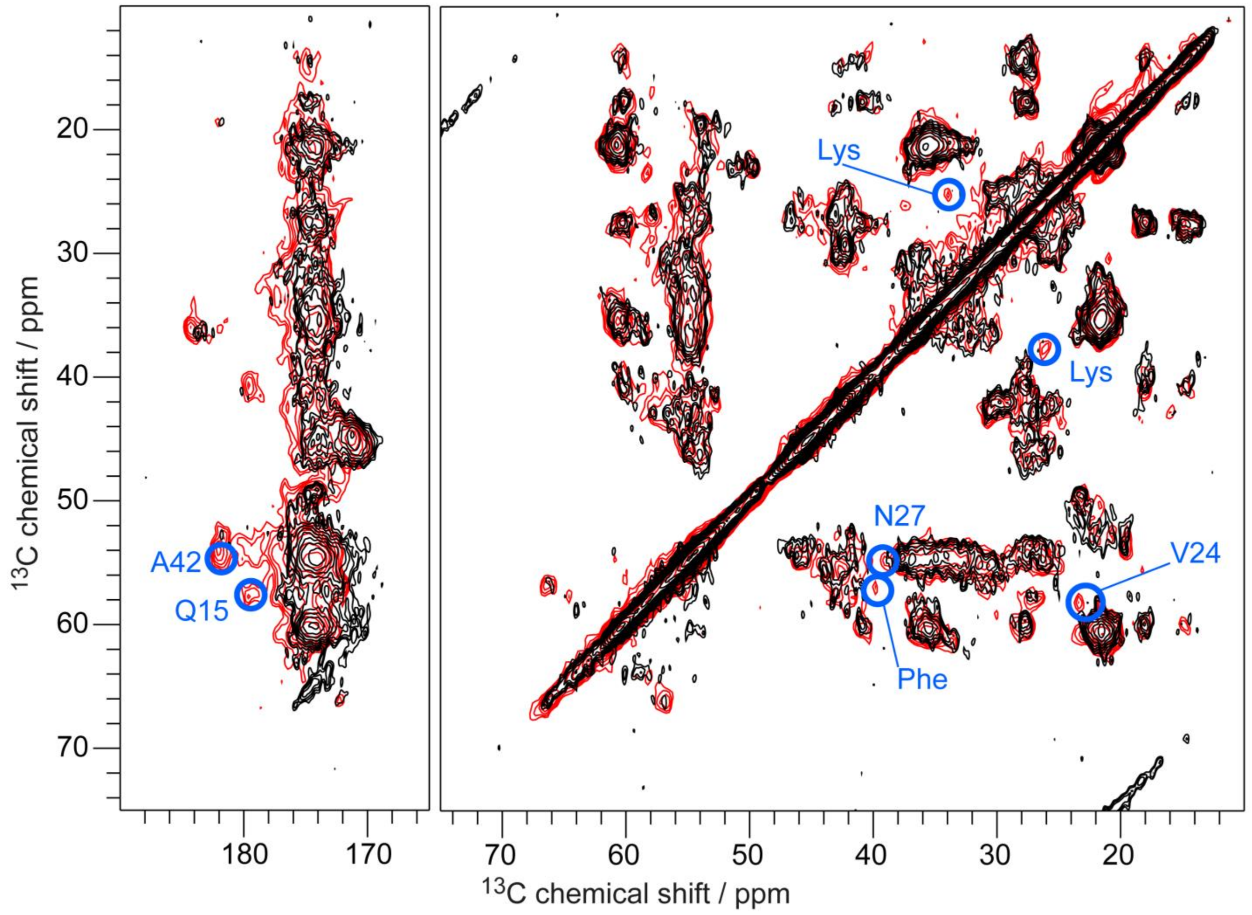

**Supplementary Fig. 19** PDSD spectra of huPrP(23-144)-Aβ* (red) (* species is $^{13}$C, $^{15}$N uniformly labeled) and huPrP(23-144)$_{exc}$-Aβ* (black) ($_{exc}$ huPrP is in excess; * species is $^{13}$C, $^{15}$N uniformly labeled), measured at a temperature of ≈ 0 °C, at a spinning frequency of 11 kHz and a mixing time of 50 ms at either a 600 MHz (huPrP(23-144)-Aβ*) or 800 MHz (huPrP(23-144)$_{exc}$-Aβ*) spectrometer. Blue circles highlight differences between the two samples.

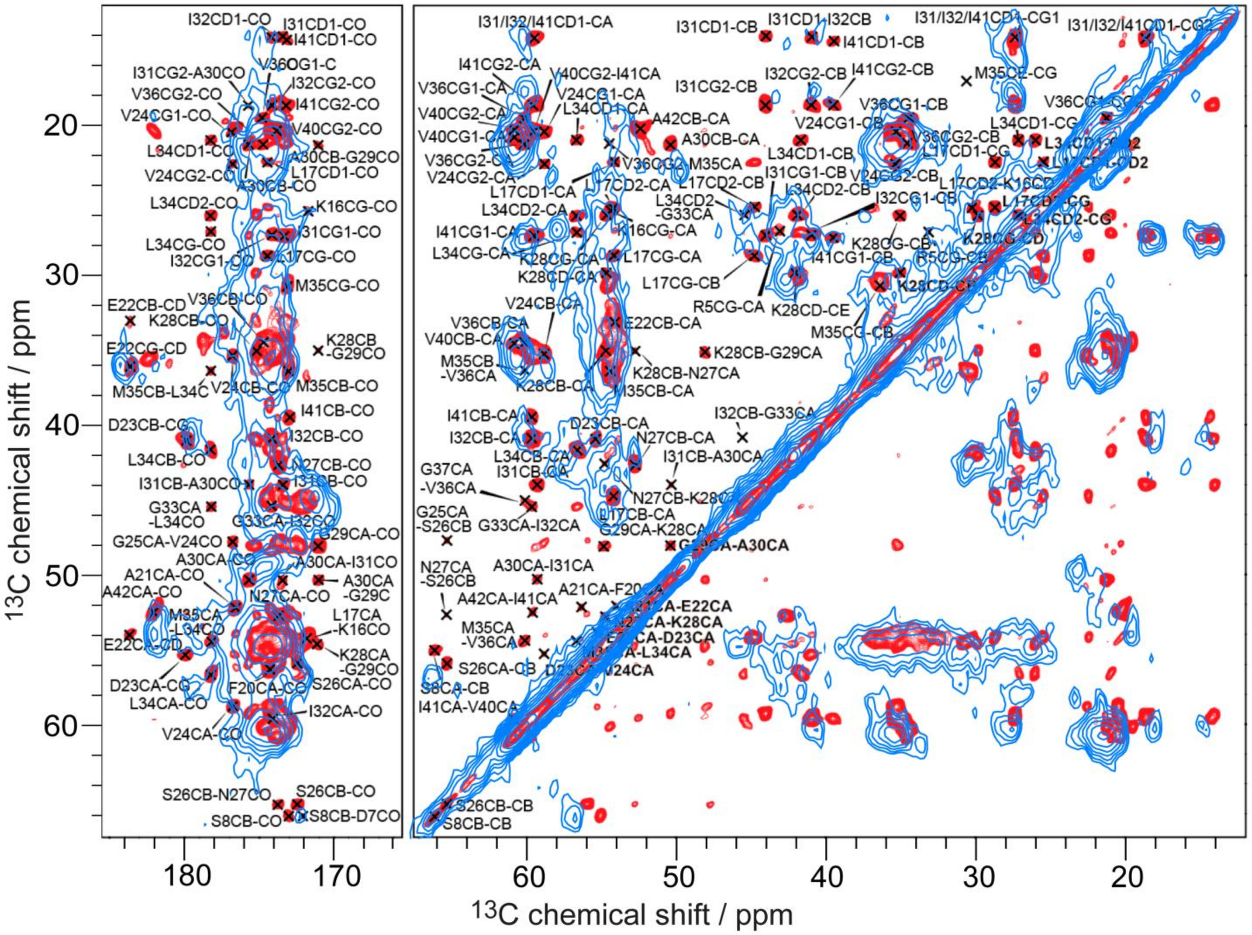

**Supplementary Fig. 20** Overlay of a PDSD spectrum (blue) of huPrP(23-144)-Aβ* (* species is $^{13}$C, $^{15}$N uniformly labeled), measured at a temperature of ≈ 0 °C, a spinning frequency of 11 kHz and a mixing time of 50 ms (same spectrum as in **Fig. 4**), and a DARR spectrum (red) of the Aβ(1-42) fibril at pH 7.4 of Colvin et al.[10], measured at a $\omega(^1H)/2\pi$ field strength of 800 MHz, a temperature of 277 K, a spinning frequency of 20 kHz and a mixing time of 80 ms. There is substantial signal overlap, but for the Aβ(1-42) fibril line widths are much smaller. Picture adapted with permission of ref. [10].

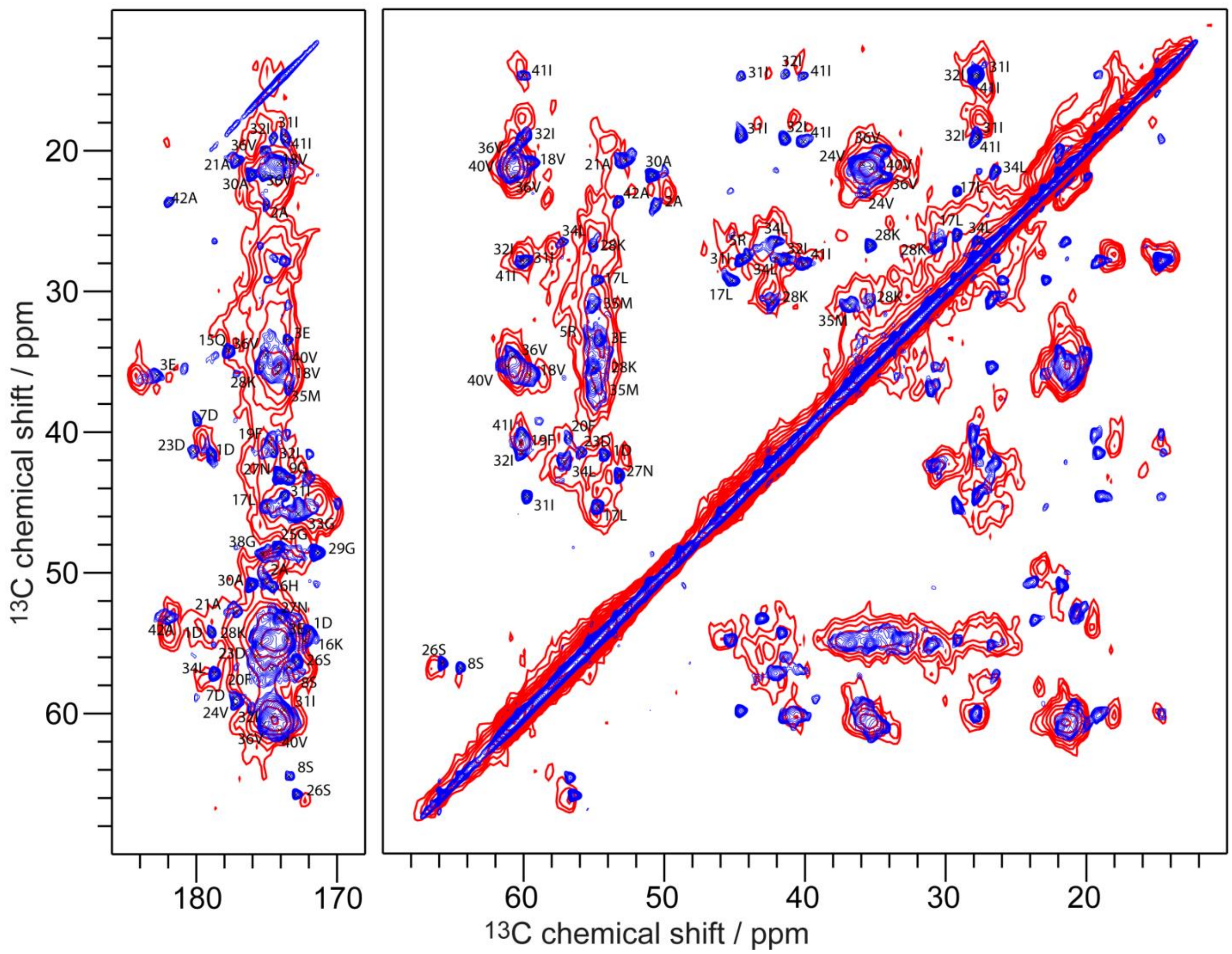

**Supplementary Fig. 21** Overlay of a PDSD spectrum (red) of huPrP(23-144)-Aβ* (* species is $^{13}$C, $^{15}$N uniformly labeled), measured at a temperature of ≈ 0 °C, a spinning frequency of 11 kHz and a mixing time of 50 ms (same spectrum as in **Fig. 4**), and a DARR spectrum (blue) of the Aβ(1-42) fibril at pH 7.4 of Ravotti et al.[11], measured at a magnetic field strength of 20.0 T, a spinning frequency of 17 kHz and a mixing time of 20 ms. There is substantial signal overlap, but for the Aβ(1-42) fibril line widths are much smaller. Picture adapted with permission of ref. [11].

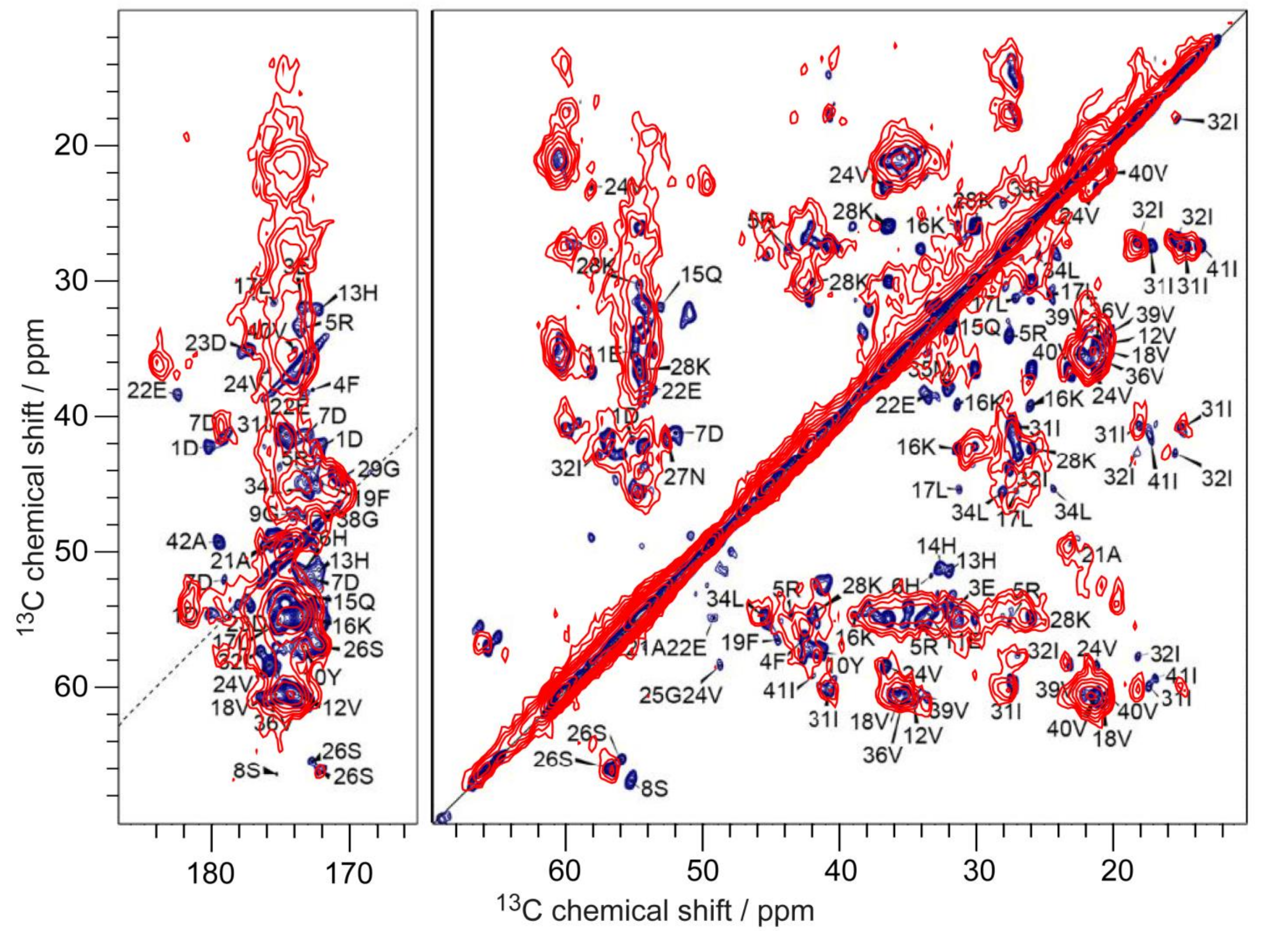

**Supplementary Fig. 22** Overlay of two PDSD spectra of huPrP(23-144)-Aβ* (* species is $^{13}$C, $^{15}$N uniformly labeled), measured at a temperature of ≈ 0 °C, a spinning frequency of 11 kHz and a mixing time of 50 ms (red, same spectrum as in **Fig. 4**), and of the Aβ(1-42) fibril at pH 2 of Gremer et al. (blue)[9], measured at a magnetic field strength of 18.8 T, a temperature of 0±5 °C, a spinning frequency of 12.5 kHz and a mixing time of 20 ms. There is substantial signal overlap, but for the Aβ(1-42) fibril line widths are much smaller. Picture adapted with permission of ref. [9].

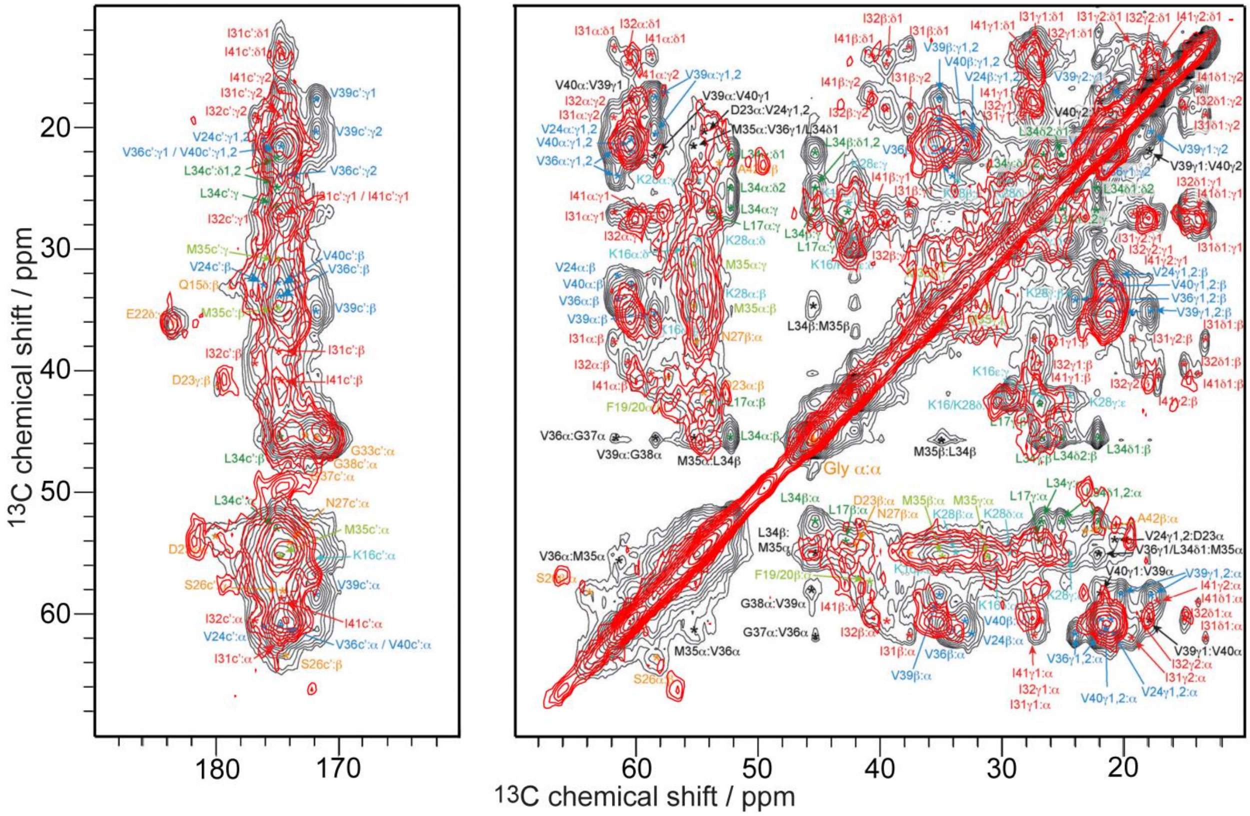

**Supplementary Fig. 23** Overlay of a PDSD spectrum (red) of huPrP(23-144)-Aβ* (* species is $^{13}$C, $^{15}$N uniformly labeled), measured at a temperature of ≈ 0 °C, a spinning frequency of 11 kHz and a mixing time of 50 ms (same spectrum as in **Fig. 4**), and a DARR spectrum (black) of a hexameric Aβ42cc protofibril (mutated at A21C, A30C) of Lendel et al.[12], measured at a $\omega(^{1}H)/2\pi$ field strength of 700 MHz, a temperature of ≈ 4 °C, a spinning frequency of 12 kHz and a mixing time of 50 ms. There is some signal overlap, although some peaks are not existing in the other Aβ species, but line widths are almost similar. Picture adapted with permission of ref. [12].

**Supplementary Table 1** Chemical shifts of unambiguously assigned resonances in huPrP(23-144)-Aβ* (* species is $^{13}C$, $^{15}N$ uniformly labeled)

| # | Amino acid | N (ppm) | CO (ppm) | CA (ppm) | CB (ppm) | CG1 (ppm) | CG2 (ppm) | CD1 (ppm) | CD2 (ppm) | CE1 (ppm) | CE2 (ppm) | CZ (ppm) | ND2 (ppm) | NE1 (ppm) | NE2 (ppm) | NZ (ppm) |
|---|---|---|---|---|---|---|---|---|---|---|---|---|---|---|---|---|
| 1 | D | | | | | | | | | | | | | | | |
| 2 | A | | | | | | | | | | | | | | | |
| 3 | E | | | | | | | | | | | | | | | |
| 4 | F | | | | | | | | | | | | | | | |
| 5 | R | 122.4 | 174.4 | 54.5 | 30.3 | 28.1 | | 42.2 | | | | - | | - | | |
| | | 126.7 | - | 53.3 | 30.9 | 25.0 | | 43.7 | | | | 165.0 | | 67.7 | | |
| 6 | H | - | - | - | - | - | | | - | - | | | | | - | |
| 7 | D | | | | | | | | | | | | | | | |
| 8 | S | | | | | | | | | | | | | | | |
| 9 | G | | | | | | | | | | | | | | | |
| 10 | Y | | | | | | | | | | | | | | | |
| 11 | E | | | | | | | | | | | | | | | |
| 12 | V | | | | | | | | | | | | | | | |
| 13 | H | - | - | - | - | - | | | - | - | | | | | - | |
| 14 | H | - | - | - | - | - | | | - | - | | | | | - | |
| 15 | Q | 119.6 | 175.0 | 57.4 | 30.5 | 32.2 | | 179.9 | | | | | | | 124.3 | |
| 16 | K | 124.7 | 174.0 | 54.8 | 35.3 | 25.9 | | 28.4 | | 42.1 | | | | | | 34.8 |
| 17 | L | 129.5 | 174.6 | 54.3 | 46.5 | 27.2 | | 24.8 | 24.8 | | | | | | | |
| 18 | V | 117.5 | 174.6 | 60.5 | 35.5 | 21.2 | 21.2 | | | | | | | | | |
| 19 | F | 122.7 | 177.0 | 56.3 | 41.0 | 137.1 | | 131.7 | 131.7 | 130.8 | 130.8 | 129.0 | | | | |
| 20 | F | 126.6 | 172.2 | 56.4 | 43.5 | 137.3 | | 131.8 | 131.8 | 129.9 | 129.9 | 127.5 | | | | |
| 21 | A | | | | | | | | | | | | | | | |
| 22 | E | | | | | | | | | | | | | | | |
| 23 | D | 123.3 | 174.6 | 54.6 | 41.6 | 178.9 | | | | | | | | | | |
| 24 | V | 122.1 | 176.6 | 58.2 | 36.8 | 23.2 | 21.6 | | | | | | | | | |
| 25 | G | 113.6 | 171.3 | 45.8 | | | | | | | | | | | | |
| 26 | S | 119.5 | 172.2 | 56.6 | 66.2 | | | | | | | | | | | |
| 27 | N | 116.4 | 174.9 | 53.0 | 43.2 | 179.1 | | | | | | | 114.3 | | | |
| | | 129.9 | 174.1 | 54.8 | 38.5 | 177.1 | | | | | | | 124.8 | | | |
| 28 | K | 119.1 | 176.8 | 54.4 | 33.7 | 25.1 | | 28.8 | | 42.2 | | | | | | 32.9 |
| 29 | G | 116.7 | 172.6 | 49.2 | | | | | | | | | | | | |
| 30 | A | 124.4 | 175.1 | 49.5 | 22.9 | | | | | | | | | | | |

| | | | | | | | | | | |
|---|---|---|---|---|---|---|---|---|---|---|
| 31 | I | 124.1 | 173.9 | 57.7 | 43.0 | 27.0 | 18.1 | 15.4 | | |
| 32 | I | 126.9 | 174.7 | 60.2 | 40.7 | 27.4 | 17.9 | 14.7 | | |
| 33 | G | 119.1 | 173.8 | 48.6 | | | | | | |
| 34 | L | | | | | | | | | |
| 35 | M | - | - | - | - | - | | | | - |
| 36 | V | | | | | | | | | |
| 37 | G | | | | | | | | | |
| 38 | G | | | | | | | | | |
| 39 | V | | | | | | | | | |
| 40 | V | 120.6 | 175.0 | 60.6 | 32.0 | 21.6 | 21.6 | | | |
| 41 | I | 122.3 | 175.3 | 60.2 | 39.4 | 27.5 | 17.9 | 13.9 | | |
| 42 | A | 117.3 | 182.1 | 53.9 | 19.4 | | | | | |

**Supplementary Table 2** Overview of the performed solid-state NMR measurements

| experiment | description | dimen-sions | transfer | purpose | samples |
|---|---|---|---|---|---|
| INEPT[13] | insensitive nuclei enhanced by polarization transfer | 1D | $^{1}$H to $^{13}$C | to selectively excite mobile regions | huPrP(23-144)*-Aβ<br>huPrP(23-230)*-Aβ<br>huPrP(23-144)-Aβ*<br>huPrP(23-144)$_{exc}$-Aβ* |
| CP | cross polarization | 1D | $^{1}$H to $^{13}$C | to excite rigid parts | huPrP(23-144)*-Aβ<br>huPrP(23-230)*-Aβ<br>huPrP(23-144)-Aβ*<br>huPrP(23-144)$_{exc}$-Aβ* |
| | | | $^{1}$H to $^{15}$N | to excite rigid parts | huPrP(23-144)*-Aβ<br>huPrP(23-144)-Aβ*<br>huPrP(23-144)$_{exc}$-Aβ* |
| PDSD[14] | proton driven spin diffusion | 2D | $^{13}$C to $^{13}$C | to excite all correlations | huPrP(23-144)*-Aβ<br>huPrP(23-230)*-Aβ<br>huPrP(23-144)-Aβ*<br>huPrP(23-144)$_{exc}$-Aβ* |
| DARR | dipolar assisted rotational resonance | 2D | $^{13}$C to $^{13}$C | to excite all correlations | huPrP(23-144)-Aβ* |
| DQ-SPC5[15] | double quantum excitation with SPC5-recoupling | 2D | $^{13}$C to $^{13}$C | to excite only one-bond correlations | huPrP(23-144)*-Aβ<br>huPrP(23-144)-Aβ* |
| NCA[16] | with SPECIFIC-CP | 2D | $^{15}$N to $^{13}$Cα | frequency selective polarization transfers from $^{15}$N to $^{13}$Cα | huPrP(23-144)*-Aβ<br>huPrP(23-144)-Aβ* |
| NCACX[16] | with SPECIFIC-CP | 2D and 3D | $^{15}$N to $^{13}$Cα to $^{13}$C$_{sidechain}$ | frequency selective polarization transfers from $^{15}$N to $^{13}$Cα, subsequent DARR mixing to distribute magnetization to side chains | huPrP(23-144)-Aβ* |
| NCOCX[16] | with SPECIFIC-CP | 3D | $^{15}$N to $^{13}$CO to $^{13}$C$_{sidechain}$ | frequency selective polarization transfers from $^{15}$N to $^{13}$CO, subsequent DARR mixing to distribute magnetization to side chains | huPrP(23-144)-Aβ* |

**Supplementary Table 3** Experimental parameters for solid-state NMR measurements of sample huPrP(23-144)*-Aβ (* species is $^{13}C$, $^{15}N$ uniformly labeled)

| **Sample huPrP(23-144)*-Aβ** | | | | | | |
|---|---|---|---|---|---|---|
| | HC CP | HN CP | INEPT | PDSD | SPC5_2 | NCA |
| | 1D | 1D | 1D | 2D | 2D | 2D |
| Mixing time (ms) | - | - | - | 1) 20<br>2) 30<br>3) 50<br>4) 100<br>5) 200<br>6) 200 $R^2$ | - | - |
| $^{1}H$ frequency (MHz) | 600 | 600 | 600 | 600 | 600 | 600 |
| MAS (kHz) | 11 | 11 | 4 | 1) & 2) & 3) & 4) & 5) 11<br>6) 9.375 | 8 | 11 |
| VT gas temperature (°C) | -10 | -16 | RT | 1) & 3) & 4) -10<br>2) & 5) & 6) -16 | -16 | -16 |
| Transfer 1 | HC CP | HN CP | INEPT | HC CP | HC CP | HN CP |
| Carrier (ppm) | 70.525 | 119.979 | 70.525 | 1) & 3) & 4) 70.525<br>2) & 5) & 6) 90.046 | 70.525 | 119.973 |
| Duration of 1st transfer (μs) | 200 | 600 | - | 1) & 3) & 4) 200<br>2) & 5) & 6) 350 | 300 | 600 |
| Transfer 2 | - | - | - | - | SPC5 | N-CA CP |
| Carrier (ppm) | - | - | - | - | 60.579 | 57.058 |
| Duration of 2nd transfer (μs) | - | - | - | - | - | 1800 |
| $^{13}C$ rf field (kHz) | - | - | - | - | - | 8.205 |
| $^{15}N$ rf field (kHz) | - | - | - | - | - | 18.803 |
| $t_1$ increments | 454 | 136 | 454 | 1) & 3) & 4) 454<br>2) & 5) & 6) 379 | 454 | 366 |
| $t_1$ spectral width (kHz) | 37.879 | 13.587 | 37.879 | 37.879 | 37.879 | 30.488 |
| $t_2$ increments | - | - | - | 1) & 5) 180<br>2) 260<br>3) & 4) 128<br>6) 190 | 240 | 22 |
| $t_2$ spectral width (kHz) | - | - | - | 1) & 3) & 4) 29.996<br>2) & 5) 33.003<br>6) 37.700 | 40.000 | 3.600 |
| Number of scans | 256 | 2000 | 256 | 1) 128<br>2) & 3) & 4) 320 | 592 | 1600 |

| | | | | | | |
|---|---|---|---|---|---|---|
| | | | | 5) 216<br>6) 568 | | |
| Duration (h) | 0.15 | 1.7 | 0.15 | 1) 26.25<br>2) 95<br>3) 47.25<br>4) 48.5<br>5) 43.75<br>6) 133.5 | 160.5 | 40 |

**Supplementary Table 4** Experimental parameters for solid-state NMR measurements of sample huPrP(23-230)*-Aβ (* species is $^{13}C$, $^{15}N$ uniformly labeled)

| **Sample huPrP(23-230)*-Aβ** | | | |
|---|---|---|---|
| | HC CP | INEPT | PDSD |
| | 1D | 1D | 2D |
| Mixing time (ms) | - | - | 1) 30<br>2) 50 |
| $^{1}H$ frequency (MHz) | 600 | 600 | 600 |
| MAS (kHz) | 11 | 11 | 11 |
| VT gas temperature (°C) | -20 | 20 | 1) -10<br>2) 0 |
| Transfer 1 | HC CP | INEPT | HC CP |
| Carrier (ppm) | 79.967 | 79.967 | 79.967 |
| Duration of 1$^{st}$ transfer (μs) | 500 | - | 200 |
| $t_1$ increments | 750 | 750 | 750 |
| $t_1$ spectral width (kHz) | 62.500 | 62.500 | 62.500 |
| $t_2$ increments | - | - | 1) 140<br>2) 134 |
| $t_2$ spectral width (kHz) | - | - | 36.199 |
| Number of scans | 512 | 5659 | 1) 1267<br>2) 272 |
| Duration (h) | 0.29 | 3.25 | 1) 201.75<br>2) 42 |

**Supplementary Table 5** Experimental parameters for solid-state NMR measurements of sample huPrP(23-144)-Aβ* (* species is $^{13}$C, $^{15}$N uniformly labeled)

| **Sample huPrP(23-144)-Aβ*** | | | | | | | | | | |
|---|---|---|---|---|---|---|---|---|---|---|
| | HC CP | HN CP | INEPT | PDSD | DARR | SPC5_2 | NCA | NCACX DARR | NCACX DARR | NCOCX DARR |
| | 1D | 1D | 1D | 2D | 2D | 2D | 2D | 2D | 3D | 3D |
| Mixing time (ms) | - | - | - | 1) 10<br>2) 30<br>3) 50<br>4) 50<br>5) 50 R²<br>6) 100<br>7) 200<br>8) 200 R²<br>9) 300 | 30 | - | - | 60 | 60 | 70 |
| $^{1}$H frequency (MHz) | 600 | 600 | 600 | 1) & 2) & 3) & 6) & 7) & 8) & 9) 600<br>4) & 5) 800 | 600 | 600 | 600 | 600 | 600 | 600 |
| MAS (kHz) | 11 | 11 | 8 | 1) & 2) & 3) & 4) & 6) & 7) & 9) 11<br>5) 12.5<br>8) 9.375 | 11 | 8 | 11 | 11 | 11 | 11 |
| VT gas temperature (°C) | -10 | -10 | 10 | -10 | -10 | -10 | -10 | -10 | -10 | -10 |
| Transfer 1 | HC CP | HN CP | INEPT | HC CP | HC CP | HC CP | HN CP | HN CP | HN CP | HN CP |
| Carrier (ppm) | 70.978 | 120.159 | 70.978 | 1) & 2) & 3) & 6) & 7) & 8) & 9) 70.978<br>4) & 5) 70.307 | 70.978 | 1) 70.525<br>2) 70.153 | 119.979 | 119.979 | 119.979 | 119.979 |
| Duration of 1st transfer (μs) | 400 | 500 | - | 400 | 400 | 1) 400<br>2) 500 | 900 | 900 | 900 | 900 |
| Transfer 2 | - | - | - | - | - | SPC5 | N-CA CP | N-CA CP | N-CA CP | N-CO CP |
| Carrier (ppm) | - | - | - | - | - | 1) 67.210 | 57.057 | 57.057 | 57.057 | 174.950 |

| | | | | | | | | | | |
|---|---|---|---|---|---|---|---|---|---|---|
| | | | | | | 2) 23.735 | | | | |
| Duration of $2^{nd}$ transfer (μs) | - | - | - | - | - | - | 1600 | 1600 | 1600 | 1800 |
| $^{13}C$ rf field (kHz) | - | - | - | - | - | - | 17.806 | 17.806-<br>18.824 | 18.824-<br>19.078 | 38.156 |
| $^{15}N$ rf field (kHz) | - | - | - | - | - | - | 28.490 | 28.490-<br>29.304 | 28.490-<br>29.304 | 27.676 |
| $t_1$ increments | 454 | 181 | 454 | 1) & 2) & 3) & 6) & 7) & 8) & 9) 454<br>4) & 5) 1250 | 454 | 1) 454<br>2) 536 | 366 | 536 | 402 | 536 |
| $t_1$ spectral width (kHz) | 37.879 | 18.116 | 37.879 | 1) & 2) & 3) & 6) & 7) & 8) & 9) 37.879<br>4) & 5) 104.167 | 37.879 | 1) 37.879<br>2) 44.643 | 30.488 | 44.643 | 44.643 | 59.524 |
| $t_2$ increments | - | - | - | 1) & 2) & 3) & 6) & 7) & 9) 200<br>4) & 5) 264<br>8) 225 | 200 | 1) & 2) 240 | 22 | 22 | 18 | 12 |
| $t_2$ spectral width (kHz) | - | - | - | 1) & 2) & 3) & 6) & 7) & 9) 33.003<br>4) & 5) 44.000<br>8) 37.700 | 33.003 | 1) & 2) 40.000 | 3.600 | 3.600 | 3.600 | 2.400 |
| $t_3$ increments | - | - | - | - | - | - | - | - | 20 | 13 |
| $t_3$ spectral width (kHz) | - | - | - | - | - | - | - | - | 5.100 | 3.300 |
| Number of scans | 128 | 2000 | 256 | 1) 304<br>2) & 3) & 5) & 7) 288<br>4) 224<br>6) 368<br>8) 320<br>9) 352 | 288 | 1) 320<br>2) 496 | 864 | 1696 | 736 | 1072 |
| Duration (h) | 0.08 | 1.75 | 0.15 | 1) 69<br>2) 66 | 66 | 1) 86.75<br>2) 133.25 | 21.25 | 159.25 | 612 | 387.75 |

| | |
|---|---|
| | 3) & 4) 66.5<br>5) & 6) 87<br>7) 71.25<br>8) 89<br>9) 91 |

**Supplementary Table 6** Experimental parameters for solid-state NMR measurements of sample huPrP(23-144)$_{exc}$-Aβ* (* species is $^{13}$C, $^{15}$N uniformly labeled)

| **Sample huPrP(23-144)$_{exc}$-Aβ*** | | | | |
|---|---|---|---|---|
| | HC CP | HN CP | INEPT | PDSD |
| | 1D | 1D | 1D | 2D |
| Mixing time (ms) | - | - | - | 1) 50<br>2) 50 (after 4 months) |
| $^{1}$H frequency (MHz) | 800 | 800 | 800 | 800 |
| MAS (kHz) | 5 | 11 | 8 | 11 |
| VT gas temperature (°C) | -3 | -10 | 10 | -10 |
| Transfer 1 | HC CP | HN CP | INEPT | HC CP |
| Carrier (ppm) | 70.183 | 124.191 | 70.000 | 70.183 |
| Duration of 1$^{st}$transfer (µs) | 600 | 600 | - | 600 |
| $t_1$ increments (half of STATES-TPPI) | 778 | 949 | 778 | 778 |
| $t_1$ spectral width (kHz) | 52.083 | 59.524 | 52.083 | 52.083 |
| $t_2$ increments (half of STATES-TPPI) | - | - | - | 256 |
| $t_2$ spectral width (kHz) | - | - | - | 50.505 |
| Number of scans | 256 | 2048 | 1024 | 432 |
| Duration (h) | 0.15 | 3 | 0.5 | 127.75 |

**Supplementary Table 7** Overview of the performed solution NMR measurements

| experiment | description | dimensions | transfer | purpose | Samples |
|---|---|---|---|---|---|
| HSQC | heteronuclear single-quantum coherence | 2D | $^{1}$H to $^{15}$N and back | amide groups | huPrP(23-144) monomers |
| HNCO | triple resonance | 3D | $^{1}$H to $^{15}$N to $^{13}$CO and back | resonance assignment | huPrP(23-144) monomers |
| HNCACB | triple resonance | 3D | $^{1}$H to $^{15}$N to $^{13}$Cα to $^{13}$Cβ and back | resonance assignment | huPrP(23-144) monomers |
| BEST-TROSY-(H)N(COCA)NH | triple resonance | 3D | $^{1}$H to $^{15}$N to $^{13}$CO to $^{13}$Cα to $^{15}$N to $^{1}$H | resonance assignment | huPrP(23-144) monomers |
| TOCSY | total correlated spectroscopy | 2D | $^{13}$C to $^{13}$C | to excite all correlations | huPrP(23-144) monomers Aβ monomers |

1 Zahn, R. *et al.* NMR solution structure of the human prion protein. *Proc. Natl. Acad. Sci.* **97**, 145-150, doi:10.1073/pnas.97.1.145 (2000).

2 Shen, Y. & Bax, A. Protein backbone and sidechain torsion angles predicted from NMR chemical shifts using artificial neural networks. *J. Biomol. NMR* **56**, 227-241, doi:10.1007/s10858-013-9741-y (2013).

3 Brener, O. *et al.* QIAD assay for quantitating a compound’s efficacy in elimination of toxic Aβ oligomers. *Sci. Rep.* **5**, 13222-13234, doi:10.1038/srep13222 (2015).

4 Rösener, N. S. *et al.* A d-enantiomeric peptide interferes with heteroassociation of amyloid-beta oligomers and prion protein. *J. Biol. Chem.* **293**, 15748-15764, doi:10.1074/jbc.RA118.003116 (2018).

5 Theint, T., Nadaud, P. S., Surewicz, K., Surewicz, W. K. & Jaroniec, C. P. 13C and 15N chemical shift assignments of mammalian Y145Stop prion protein amyloid fibrils. *Biomol. NMR Assign.* **11**, 75-80, doi:10.1007/s12104-016-9723-6 (2017).

6 Han, B., Liu, Y., Ginzinger, S. W. & Wishart, D. S. SHIFTX2: significantly improved protein chemical shift prediction. *J. Biomol. NMR* **50**, 43-57, doi:10.1007/s10858-011-9478-4 (2011).

7 Schulte, C. F. *et al.* BioMagResBank. *Nucleic Acids Res.* **36**, D402-D408, doi:10.1093/nar/gkm957 (2007).

8 Calzolai, L. & Zahn, R. Influence of pH on NMR Structure and Stability of the Human Prion Protein Globular Domain. *J. Biol. Chem.* **278**, 35592-35596, doi:10.1074/jbc.M303005200 (2003).

9 Gremer, L. *et al.* Fibril structure of amyloid-beta(1-42) by cryo-electron microscopy. *Science* **358**, 116-119, doi:10.1126/science.aao2825 (2017).

10 Colvin, M. T. *et al.* High Resolution Structural Characterization of Aβ42 Amyloid Fibrils by Magic Angle Spinning NMR. *J. Am. Chem. Soc.* **137**, 7509-7518, doi:10.1021/jacs.5b03997 (2015).

11 Ravotti, F. *et al.* Solid-state NMR sequential assignment of an Amyloid-β(1–42) fibril polymorph. *Biomol. NMR Assign.* **10**, 269-276, doi:10.1007/s12104-016-9682-y (2016).

12 Lendel, C. *et al.* A Hexameric Peptide Barrel as Building Block of Amyloid-β Protofibrils. *Angew. Chem.* **126**, 12970-12974, doi:10.1002/ange.201406357 (2014).

13 Morris, G. A. & Freeman, R. Enhancement of nuclear magnetic resonance signals by polarization transfer. *J. Am. Chem. Soc.* **101**, 760-762, doi:10.1021/ja00497a058 (1979).

14 Szeverenyi, N. M., Sullivan, M. J. & Maciel, G. E. Observation of spin exchange by two-dimensional fourier transform 13C cross polarization-magic-angle spinning. *J. Magn. Reson. (1969)* **47**, 462-475, doi:10.1016/0022-2364(82)90213-X (1982).

15 Hohwy, M., Rienstra, C. M., Jaroniec, C. P. & Griffin, R. G. Fivefold symmetric homonuclear dipolar recoupling in rotating solids: Application to double quantum spectroscopy. *J. Chem. Phys.* **110**, 7983-7992, doi:10.1063/1.478702 (1999).

16 Baldus, M., Petkova, A. T., Herzfeld, J. & Griffin, R. G. Cross polarization in the tilted frame: assignment and spectral simplification in heteronuclear spin systems. *Mol. Phys.* **95**, 1197-1207 (1998).
